# CNV-Profile Regression: A New Approach for Copy Number Variant Association Analysis in Whole Genome Sequencing Data

**DOI:** 10.1101/2024.11.23.624994

**Authors:** Yaqin Si, Wenbin Lu, Shannon Holloway, Hui Wang, Albert A. Tucci, Amanda Brucker, Yuhuan Cheng, Li-San Wang, Gerard Schellenberger, Wan-Ping Lee, Jung-Ying Tzeng

## Abstract

Copy number variants (CNVs) are DNA gains or losses involving >50 base pairs. Assessing CNV effects on disease risk requires consideration of several factors. First, there are no natural definitions for CNV loci. Second, CNV effects can depend on dosage and length. Third, CNV effects can be more accurately estimated when all CNV events in a genomic region are analyzed together to assess their joint effects. We propose a new framework for association analysis that directly models an individual’s entire CNV profile within a genomic region. This framework represents an individual’s CNVs using a CNV profile curve to capture variations in CNV length and dosage and to bypass the need to predefine CNV loci. CNV effects are estimated at each genome position, making the results comparable across different studies. To jointly estimate the effects of all CNVs, we use a Lasso penalty to select CNVs associated with the trait and integrate a weighted L2-fusion penalty to encourage similar effects of adjacent CNVs when supported by the data. Simulations show that the proposed model can more effectively identify causal CNVs while maintaining false positive rates comparable to baseline methods and yield more precise effect-size estimates across different settings. When applied to CNV derived from whole genome sequencing data of the Alzheimer’s Disease Sequencing Project, the proposed methods identify additional CNVs associated with Alzheimer’s Disease (AD). These identified CNVs overlap with several known AD-risk genes and are significantly enriched by biological processes related to neuron structures and functions crucial in AD development.

## Introduction

Copy number variants (CNVs) are DNA gains or losses that can range in length from 50 base pairs (bp) to megabases (Mb).^1^ CNVs cover 5%– 12% of the genome in different populations and encompass more nucleotide base pairs than single nucleotide polymorphisms (SNPs).^2,3^ In addition to Mendelian disorders, CNVs are also associated with complex traits ranging from seemingly normal physical morphology (e.g., brain structure divergences^4^) to disorders such as Alzheimer Diseases (AD)^5–7^ and schizophrenia^8,9^. Phenomewide evaluations of whole-genome CNVs have also identified novel CNV-phenotype associations and broadened our understanding of the potential effects of CNVs on human health and diseases.^10–13^ As next-generation sequencing (NGS) becomes more accessible, whole genome sequencing (WGS) offers a thorough examination of all types of CNVs (small and large; common and rare) with comprehensive coverage and higher resolution, enabling an in-depth study of CNVs in diseases.^11,14^

One crucial step to conducting CNV association analysis is to define a “CNV locus”. Unlike SNPs, there is no natural definition of “locus” for CNVs due to breakpoints nonalignment (i.e., random CNV starting or ending positions across different individuals). CNVs spanning multiple base pairs often overlap partially between individuals, particularly with WGS data. Several CNV locus definitions have been considered in the literature and may be used in CNV analysis with WGS data. (1) Recurrent CNV (RC) focuses on CNVs that have been reported to affect clinical phenotypes in established databases and publications.^10,15–17^ (2) Probe (PB) (or breakpoint) uses probes on the array as a CNV locus.^12,18^ (3) Overlapping with density trimming (DT) first merges CNVs of different individuals into a larger “region” if CNVs overlap by *>* 1 bp, and then trims off those regions with low density of CNVs (e.g., *<* 10% of the total contributing CNVs).^19–21^ (4) Reciprocal overlapping (RO) defines a CNV locus as a region that contains a set of overlapping CNVs with sufficient (e.g., 50%) mutual overlap.^11,19^ (5) Fragment (FG) defines a CNV locus as a region between two consecutive CNV breakpoints across all individuals.^19,22^

Different locus definitions can result in different total numbers of CNV loci and different lengths of CNV loci for downstream association analysis. Even if the same definition is used, for PB, DT, RO, and FG, the derived CNV loci may still vary across different studies due to unique sample overlapping patterns in each study. Similarly, for the same study, including additional samples could also change the defined CNV loci due to new overlapping patterns observed. These factors make it difficult to compare the effect of the identified trait-associated CNVs across different studies and to combine association results using meta analysis.

Given the CNV loci defined, a common practice of CNV association analysis is to apply methods used for studying SNP associations. For example, CNVranger^23^, CoNVAQ^24^, and ParseCNV2^18^ perform Fisher’s exact test, chi-squared test, and linear or logistic regression for one CNV at a time. However, adopting SNP-based approaches tends to focus only on the effect of CNV dosages (e.g., deletion, normal copy, and duplication) and does not incorporate CNV length information. As we illustrate in the Methods section, when regressing the trait on CNV dosage, the estimated effect size is indeed an aggregated effect over the genomic region covered by the CNV locus and hence would depend on the CNV length. This introduces further complexity when comparing CNV effect sizes across various studies. CNV length also contributes to complex traits – burden analyses incorporating CNV length have demonstrated an impact of total length on complex traits.^8,13^ Compared to methods only considering dosage, incorporating both dosage and length features can further boost detecting power as shown in CNV collapsing methods, such as CCRET^9^, CKAT^25^, MCKAT^26^, and CONCUR^27^.

To accommodate the high-resolution CNV calls from WGS data and incorporate CNV length and dosage polymorphisms, in this study, we follow the idea of Brucker et al.^27^ and represent an individual’s CNV events using a piecewise constant curve called “CNV profile” curve (Figure 1a) over a genomic region (e.g., chromosome). The association is then evaluated by regressing trait values on the entire CNV profile curve without introducing CNV locus definitions while adjusting for covariates. The corresponding CNV effects are obtained at each genome position and described using the “CNV effect” curves (Figure 1b), which can be directly compared across different studies. To control model complexity and leverage CNV structural information in model estimation, we apply an L_1_ Lasso penalty to select important CNVs associated with the phenotype. Additionally, we impose an weighted L_2_-fusion penalty on the effect differences between adjacent CNVs to encourage similar effect sizes, with weights introduced to control the level of smoothing between adjacent CNVs. The penalized regression model (Figure 1c) encourages sparsity in variable selection and effect smoothness between consecutive CNV events.

**Figure 1:**
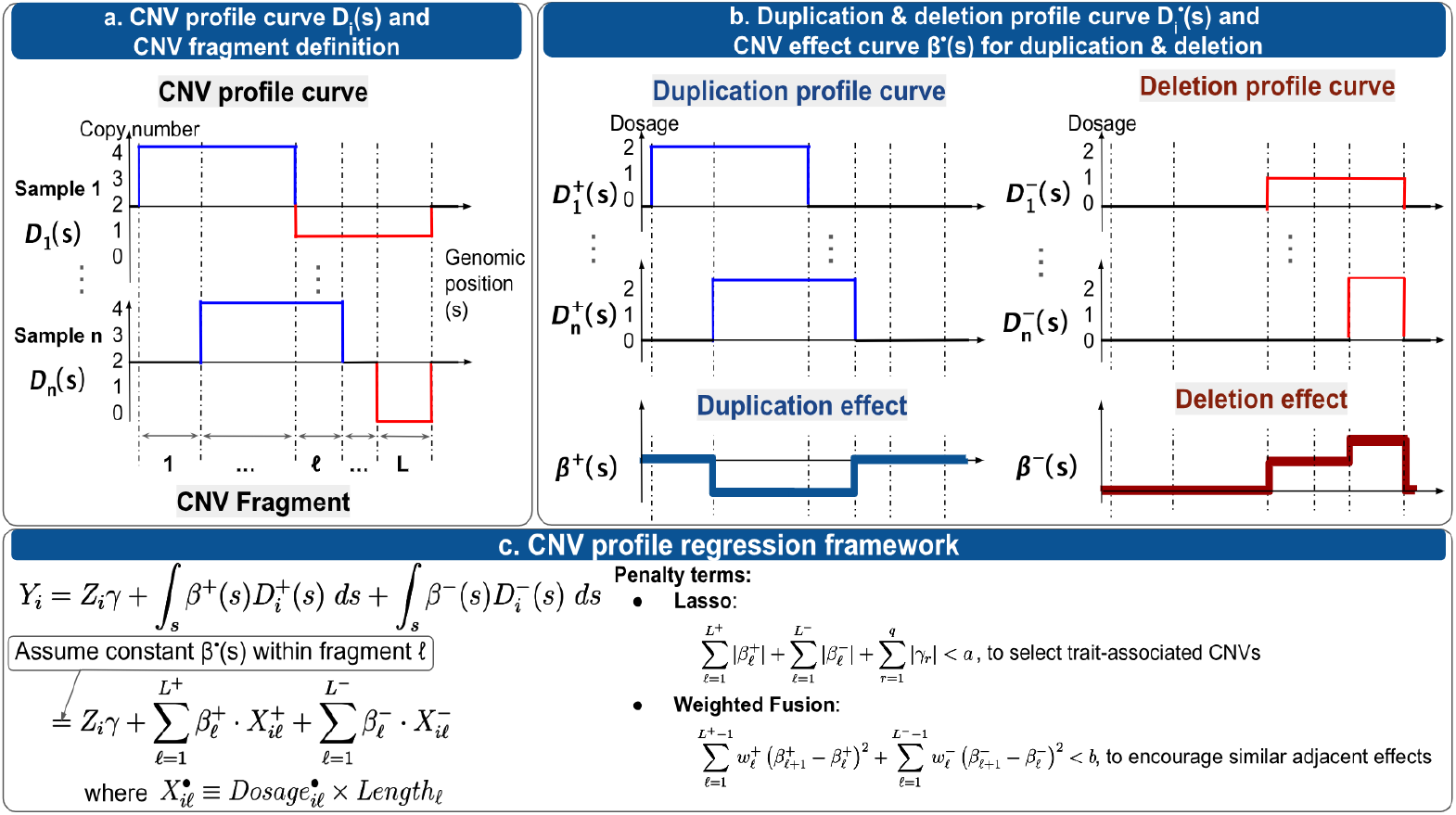
Graphical abstract. **a.** The CNV profile curve, denoted as D_i_(s), represents the CNV events of individual i across a genomic region. CNV “fragments”, one commonly used CNV locus definition, can be defined based on the breakpoints observed. **b**. The CNV profile curve can be decomposed into a duplication profile curve 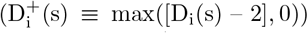 and a deletion profile curve 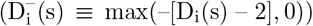. The effects of duplication/deletion on the traits are captured by the duplication/deletion effect curves. **c**. CNV profile regression evaluates the association between CNVs and trait value Y by regressing Y on the duplication and deletion profile curves, without the need to pre-define CNV loci. If assuming constant effects within a CNV fragment, the model can be reduced to model CNV effects at the fragment level, where the CNV predictor (X) is the product of CNV dosage and fragment length. The regression uses an L_1_ Lasso penalty to select trait-associated CNVs and uses a weighted L_2_ fusion penalty to encourage similar adjacent effects.

The proposed method is evaluated using simulations and real data applications. We conduct extensive simulations under various causal signal patterns and strengths to evaluate the performance of variable selection and effect estimation. We apply the proposed methods to CNVs from WGS of the Alzheimer’s Disease Sequencing Project (ADSP) to identify CNVs associated with Alzheimer’s Disease (AD). We then examine genes disrupted by the identified AD-associated CNVs, annotate them using Gene Ontology (GO) terms^28,29^, and investigate enriched biological processes to understand their relevance to AD.

## Methods

### Formulation of CNV profile regression

Consider a study consisting of n individuals. For individual i, i = 1, …, n, Y_i_ is the trait value that can be continuous or binary; Z_i_ is the q × 1 covariate vector including the intercept term; D_i_(s) is the copy number of CNV at genome position s. The plot of D_i_(s) over s, as illustrated in Figure 1a, depicts the copy number at position s for individual i and is referred to as the CNV profile curve^27^. The unit of s in the CNV curve can be set at bps or 100 bps, depending on the resolution of the CNV calling technology.

To evaluate CNV association, we can regress the trait value on the CNV profile curve D_i_(s) while adjusting for covariates Z_i_, using a generalized linear model (GLM). However, doing so constrains the effects of duplication and deletion at position s to be the same.

To allow for unconstrained effects of duplication and deletion, we define

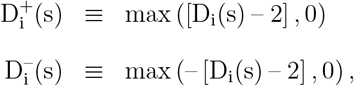

where 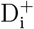 and 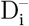 represent the duplication profile curve and deletion profile curve of individual i, respectively (e.g., Figure 1b). Here the subtraction of 2 in 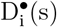 reflects the baseline copy number 2. We then propose the following model:

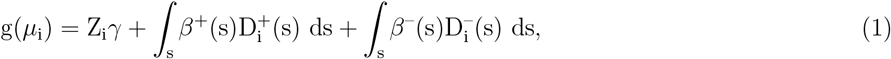

where 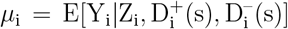 is the conditional expectation of Y_i_; g (·) is the link function with g(*µ*_i_) = *µ*_i_ for continuous traits and g(*µ*_i_) = logit(*µ*_i_) = log [*µ*_i_/(1 – *µ*_i_)] for binary traits; *γ* is a q × 1 vector for covariate coefficients including the intercept; and *β*^+^ (s) and *β*^−^ (s) are the effect coefficients for duplications and deletions at the position s, respectively.

Model (1) can be connected to the commonly used CNV fragment (FG) loci^19^ (e.g., Figure 1a). According to the fragment definition^19^, an individual i has the same dosage across all positions within a CNV fragment. That is, for fragment *ℓ, ℓ* = 1, · · ·, L, we have 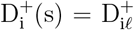 and 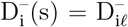 for s ∈ FG *ℓ*. Therefore, it is reasonable to assume a constant effect across position s within fragment *ℓ*, i.e., 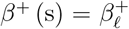 and 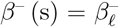 for s ∈ FG *ℓ*. Consequently, Model (1) can be simplified to model CNV effects at the level of “CNV fragment”, rather than at each unit-position s (e.g., bp). To see this, let T_*ℓ*_ be the length of fragment *ℓ*. Then under the assumption of constant effects within a fragment, Model (1) can be expressed as follows.

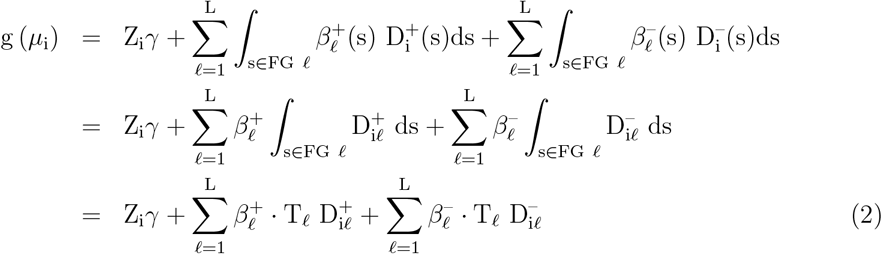

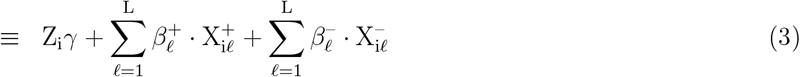

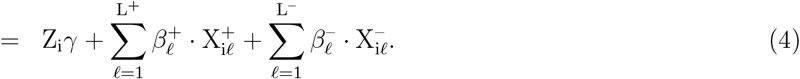

Equations (2) and (3) follow because 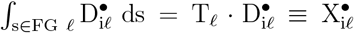, where the new variable 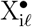 incorporates both the length and dosage information of a duplication/deletion fragment. The corresponding coefficient 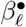 is the CNV duplication/deletion effect of fragment *ℓ*. Because not every fragment contains both duplications and deletions, we can further rewrite Equation (3) to Equation (4), where L^+^ (and L^−^) denote the total number of fragments with observed duplication (and deletion) events and L^•^ ≤ L. Such re-expression allows us to focus only on fragments with observed events of interest.

To gain more insight about 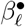,consider a commonly adopted alternative CNV model that regresses trait values on CNV dosage of a fragment:

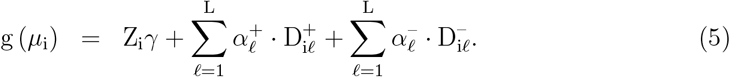

Comparing Equations (2) and (5), we see that coefficient 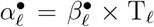, indicating that 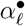 is the “aggregate” CNV effect across all s’s in fragment *ℓ*. Because 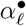 depends on the length of fragment *ℓ* and fragment lengths can vary across different studies, CNV effect via 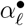 is not directly comparable across different studies. In contrast, CNV effect size 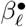 is directly comparable across studies because it represents the CNV effect size at position s for s within fragment *ℓ*.

### Implementation of CNV profile regression

One issue of working with CNV fragment loci is that a continuous CNV event can be artificially split into multiple consecutive CNV fragments. Although Model (4) allows fragment-specific effects 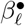, adjacent CNV fragments are expected to have similar effect coefficients, i.e., 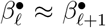 when fragments *ℓ* and *ℓ* + 1 are adjacent. The similar effect sizes arise because adjacent CNV fragments are likely to disrupt the same functional elements in the genome. To address the artificial partitioning of continuous CNV events, we impose a weighted L_2_-fusion penalty^30^ on the adjacent CNV fragments to encourage similar effect coefficients 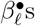 between two consecutive fragments. Additionally, we also impose an L_1_ Lasso penalty^31^ to select trait-associated CNV fragments. These penalty terms allow the CNV profile regression to estimate association effects, smooth effect sizes for adjacent CNV fragments, and select trait-associated fragments simultaneously.

To estimate regression parameters, define the design matrices of duplications and deletions as 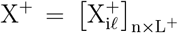 and 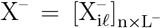, respectively, where 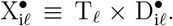 Let L^∗^ = L^+^ + L^−^ and define X_n×L_^∗^ ≡ (X^+^, X^−^), *β*_L∗ ×1_ ≡ (*β*^+⊤^, *β*^−⊤^)^⊤^, and *θ*_(L∗+q)×1_ = (*β*^⊤^, *γ*^⊤^)^⊤^. We propose to estimate parameter *θ* by

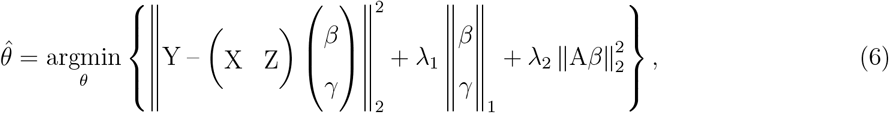

where the L_1_ penalty

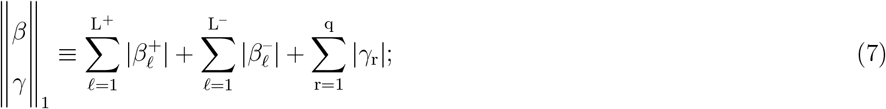

and the L_2_ penalty

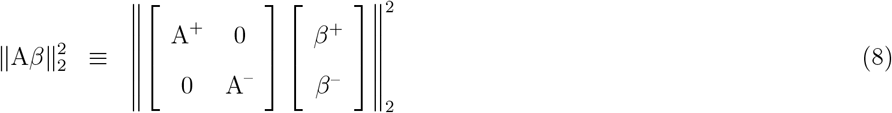

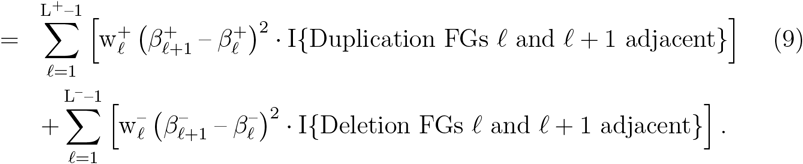

In Equation (8), A^•^ is an (L^•^ – 1) × L^•^ matrix with entry 0 except that 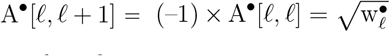.I{• FGs *ℓ* and *ℓ* + 1 adjacent}. For example, assuming that there are 7 duplication fragments in total, where the 2nd-4th fragments as well as the 6th-7th fragments are adjacent. Then the corresponding A^+^ is

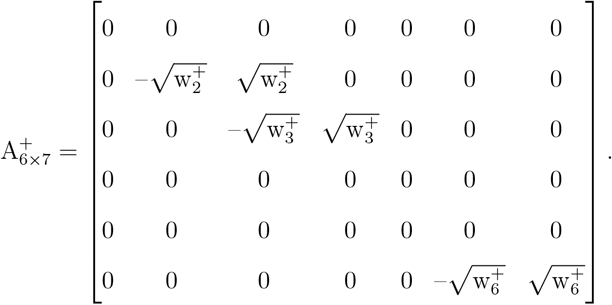

The weight term 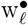 controls the magnitude of smoothing between adjacent CNV fragments *ℓ* and *ℓ* + 1. We consider two types of weights: (a) equal weight, i.e., 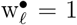 for all adjacent *ℓ*’s; and (b) correlation weight, i.e., 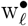 is set as the cosine similarity between CNV events in fragments *ℓ* and *ℓ* + 1. We use cosine similarity to quantify correlation because the CNV design vector 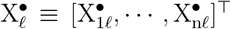 tends to be sparse and contains many 0’s, and cosine similarity can quantify similarities based on the non-zero values in common between 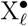 and 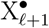. Specifically, the cosine similarity between fragments *ℓ* and *ℓ* + 1 is

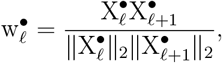

where 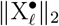 is the L_2_ norm of vector 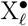,defined as 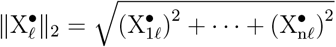. A cosine value of 0 means that the two CNV fragments are orthogonal with zero correlation. The closer the cosine value is to 1, the greater the correlation between the two CNV fragments.

The tuning parameters λ_1_ and λ_2_ control the level of penalization for sparsity (λ_1_) and smoothness (λ_2_). As λ_1_ increases, the model increases in sparsity, leading to fewer variables with non-zero coefficients. As λ_2_ increases, the coefficients for adjacent CNV fragments become more similar to each other. When λ_2_ = 0, the model reduces to a Lasso regression. We use 5-fold cross-validation (CV) and grid search on the combination of candidate λ_1_ and λ_2_ to identify the optimal tuning parameter values. The best pair of tuning parameters that minimizes the average validation loss in the 5-fold CV is used to determine the final estimate of *θ*. We describe the estimation details for the penalized CNV profile regression in Appendix A and provide the computation pseudo-code in the supplemental information.

### Simulation studies

#### Simulation designs

We use simulations to evaluate the performance of the proposed methods. We obtain CNV data from the WGS of the ADSP, and focus on the p-arm of chromosome 10 from the n = 6, 665 Non-Hispanic White (NHW) individuals.^32^ To maintain the necessary resolution without generating excessive noise, we round the position of CNV breakpoints to the nearest 25 base pair positions. We then define CNV fragments using CNVruler^19^, and use those CNV fragments with CNV event frequency *>*5% for simulations. There are a total of L = 270 CNV fragments, including L^+^ = 14 fragments with duplications and L^−^ = 256 fragments with deletions. There are no fragments with both duplications and deletions in the samples. The length and frequency of these CNV fragments are shown in **??** in the supplemental information.

Next we simulate trait value Y_i_ of individual i, i = 1, · · ·, n, based on the CNV data from ADSP. For continuous traits, we generate Y_i_ from the model: 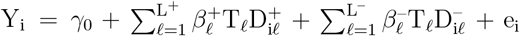, where e_i_ is generated from N(0, 1). For binary traits, we generate Y_i_ from a Bernoulli distribution with 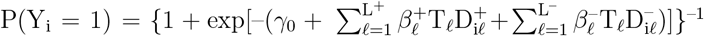. We set *γ*_0_ = –2 for both continuous and binary traits. The CNV effect 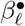 is set to 0 for “non-causal” fragments (i.e., fragments with zero effects on traits) and is determined as follows for “causal” fragments (i.e., fragments with non-zero effects on traits).

We pre-select some genomic regions as candidate causal regions to assign non-zero effects and consider three scenarios of CNV effects within these regions as summarized in Table 1: (1) causal CNV fragments are within a sub-region in the candidate regions; (2) causal CNV fragments span the entire candidate regions; and (3) causal CNV fragments are distributed across two non-adjacent sub-regions in the candidate regions. For each scenario, we consider different signal strengths: high (H), medium (M), and low (L), where 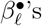 range from 0.1 to 1.5 for binary traits and from 0.1 to 0.6 for continuous traits.

**Table 1:**
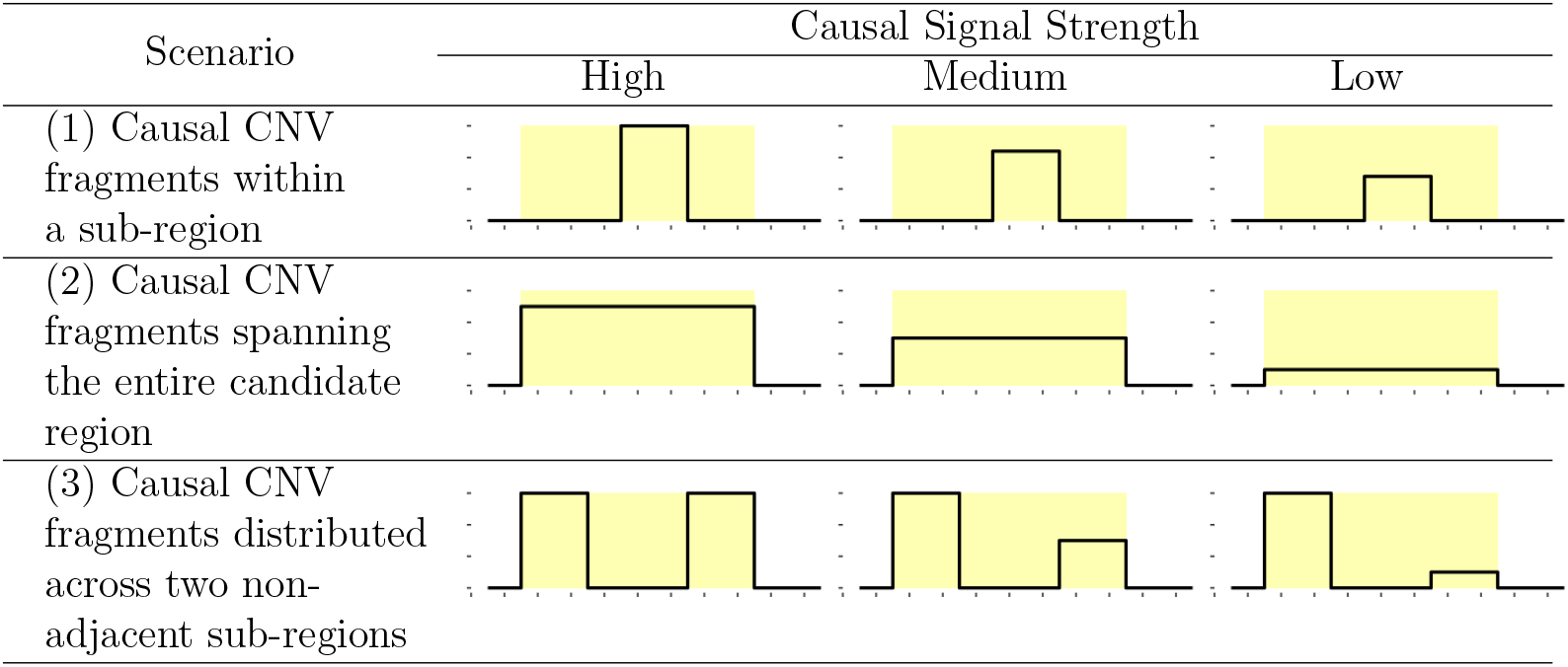
Summary of simulation settings.

#### Baseline methods and performance metrics

We evaluate the performance of the new CNV profile regression framework, i.e., smoothing with equal weight (referred to as ‘`New_eql`’) and smoothing with correlation weight (referred to as ‘`New_cor`’), and compare their performance with two frequently considered penalized methods, i.e., the standard ‘Lasso’ regression^31^ (referred to as ‘Lasso’) and the group bridge^33^ regression (referred to as ‘`gBridge`’). The `Lasso` model is equivalent to the proposed model with λ_2_ = 0 and allows us to evaluate the utility of the smoothness regularization among adjacent CNV fragments. The `gBridge` model, like the proposed methods, also accounts for the structure of consecutive CNV fragments by considering a cluster of consecutive CNV fragments as a “group”. The `gBridge` model imposes a bridge penalty on the L_1_ norms of the coefficients from the same group. That is, let g_+_ (or g_–_) indicate a group of consecutive duplication (or deletion) fragments, and let 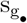 be the corresponding group size, i.e., the number o adjacent CNV fragments in group g. The group bridge penalty is 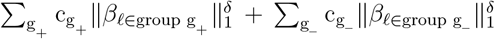 with δ = 0.5 and 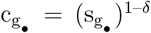 for adjusting different group sizes. Such group bridge penalty can perform variable selection at the group level and at the within-group individual variable level simultaneously.^33^ Lasso is solved with R package *glmnet* ^34^ and `gBridge` is solved using R package *grpreg* ^35^. Five-fold cross-validation (CV) is used to select the best tuning parameters, i.e., those yielding the minimum average validation loss. The selected tuning parameters are then applied to the original dataset to generate the final model and parameter estimates.

We conduct 100 simulation replications to evaluate the performance of variable selection and effect estimation. For variable selection, we compare the true positive rates (TPR), false positive rates (FPR), and the geometric mean (G-mean) of TPR and 1-FPR, which is a compositional measurement of the true positive rates and false positives rates. TPR is calculated by dividing the number of true positives (i.e., selected causal fragments) by the total number of causal fragments in each replication; then averaging this proportion over 100 replications. FPR is calculated by dividing the number of false positives (i.e., selected non-causal fragments) by the total number of non-causal fragments in each replication; then averaging this proportion over 100 replications. The G-mean is the geometric mean of TPR and (1-FPR), and is calculated by computing 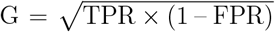 in each replication, and then averaging over 100 replications. For effect size estimation, we compute the mean squared error (MSE) of the 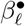 estimates as follows. First, in each replication, we compute the squared error, 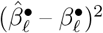,for each fragment *ℓ*, and then separately average the squared errors across causal and non-causal fragments. We then compute the mean of these average squared error over 100 replications for causal fragments and non-causal fragments, respectively. Throughout our paper, we refer to this mean of average squared error as MSE.

### ADSP analysis

Alzheimer’s disease (AD) is a progressive neurodegenerative disease and the most common cause of dementia.^36^ There are approximately 7 million Americans with AD and 32 million worldwide, making AD one of the most pressing public health issues.^36–38^ Multiple studies have highlighted the roles of CNVs in AD.^7,32,39–42^ Here we conduct CNV association analysis using CNV data obtained from ADSP WGS dataset^32^, employing the proposed and baseline methods to identify CNVs associated with AD. We focus our analysis on high-quality CNV calls generated from the GraphTyper2 caller^43^. After excluding samples with missing information on age and AD status, there are 6,665 NHW individuals (2891 controls and 3774 cases) for the CNV association analysis. As in the simulations, we round the CNV breakpoints to the nearest 25 base pair positions and define CNV fragments using CNVRuler^19^. We remove CNV fragments with an event frequency *<*5%. We conduct CNV association analysis for each chromosome arm (i.e., p-arms and q-arms of chromosomes 1 to 22), adjusting for age, sex, sequencing center, sequencing technologies, the top 10 principal components for population substructures obtained from GWAS SNPs, and the APOE genotype. For APOE genotypes, we create dummy variables for each specific genotype and use the most common “e3e3” genotype as the reference group. CNV fragments with non-zero effect size estimates are reported as AD-associated CNVs.

To understand the biological relevance of the identified CNVs with AD, we examine genes overlapping with the identified CNV fragments of each method. Specifically, we compare these identified genes with expert-curated AD-risk genes compiled by the ADSP project (https://adsp.niagads.org/gvc-top-hits-list/); we also conduct enrichment analysis with Gene Ontology (GO) annotations.^28,29,44^ First, we annotate the identified CNV fragments from each method for their effects on transcripts, proteins, and regulatory regions using the Ensembl Variant Effect Predictor (VEP)^45^ and generate a list of overlapping genes. Next, we cross-reference the overlapping genes with the current list of “AD loci with genetic evidence” and the list of “AD risk/protective causal genes”. These lists are curated by the ADSP Gene Verification Committee and only include genes that are supported by high-quality genetic evidence. For GO enrichment analysis, we convert the overlapping gene lists to Entrez IDs using the R packages *AnnotationDbi* ^46^ and *org*.*Hs*.*eg*.*db*^47^. We then input the Entrez IDs into the R package *clusterProfiler* ^48^ to identify enriched biological processes. This analysis functionally profiles the gene lists and examines the annotated terms to determine which GO terms are enriched by the genes identified by each method.^44^

Besides assessing the potential impact of the selected CNVs on AD risk by examining their biological relevance, we also conduct “null” analyses using data with permuted AD status as a negative control. We generate 50 sets of permuted outcome values and report the average of FPRs for each method.

## Results

### Simulation results

The results for binary traits under Scenario 1 (i.e., only a sub-region of consecutive CNV fragments being causal) are shown in Figure 2 (variable selection) and Figure 3 (effect estimation). Figures 2 shows that the proposed models (`New_eql` and `New_cor`) have higher TPRs than the baseline models (`Lasso` and `gBridge`); their FPRs are lower than `Lasso` but higher than `gBridge`. The results of G-mean, which provides an overall assessment by integrating both TPR and FPR, show that the proposed models have better overall variable selection performance compared to the baseline models. Figure 3 show that the proposed models yield lower MSEs for estimating causal effect sizes in causal regions. All methods show a negligible MSE (e.g., 10^−3^) for estimating the null effects. We also observe that for the proposed methods, both weighting schemes yield very similar performance in variable selection and effect estimations. This may suggest that the correlation levels among adjacent CNV fragments tend to be similar, so the correlation weight and equal weight have similar impact on smoothing.

**Figure 2:**
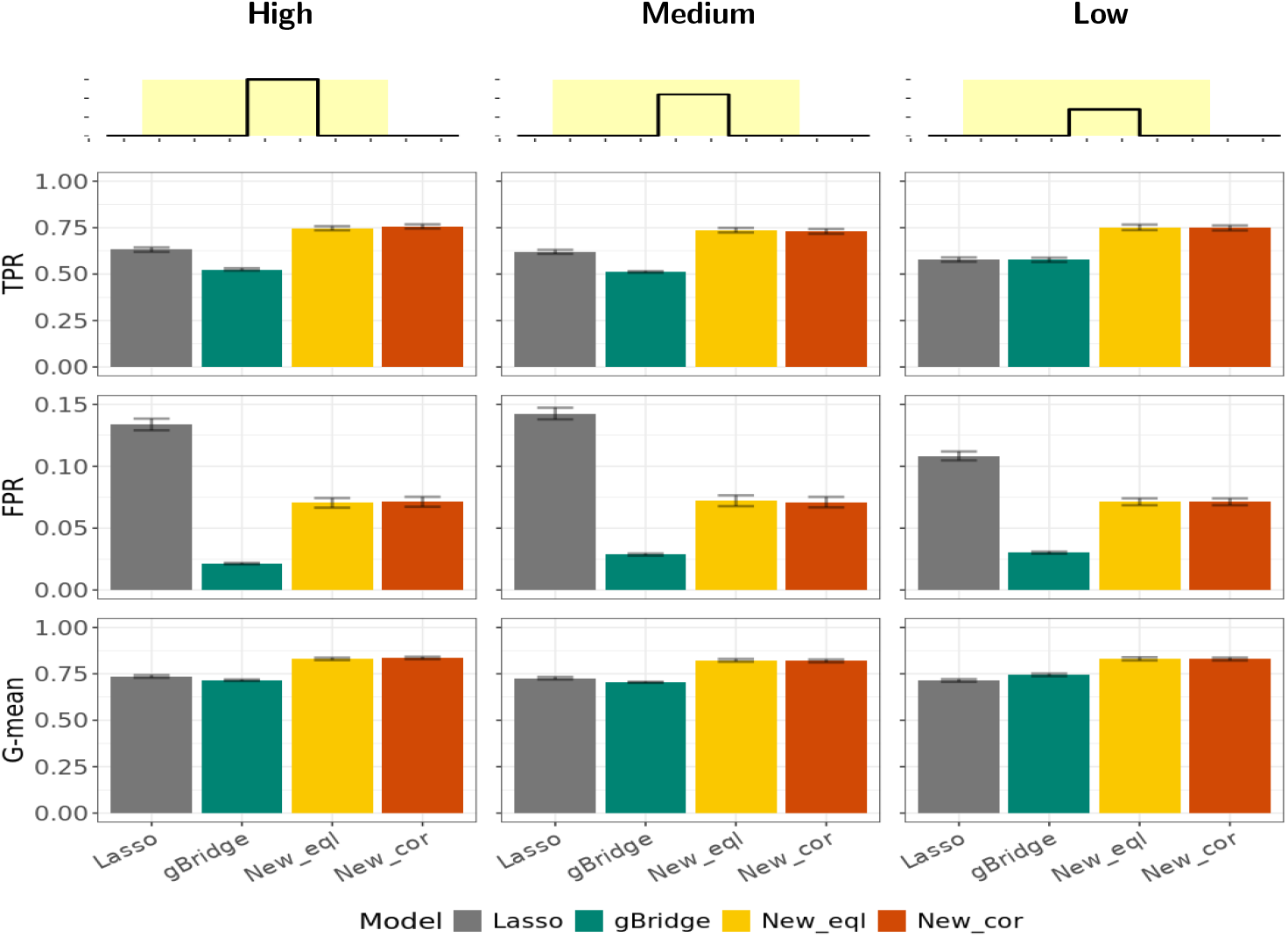
Variable selection performance evaluated by TPR, FPR, and G-mean under Simulation Scenario 1 for binary traits. The top row illustrates the design of causal signals under Scenario 1, with three levels of signal strength: high (left column), medium (middle column) to low (right column). The second to fourth rows respectively show the true positive rate (TPR), the false positive rate (FPR), and the geometric mean (G-mean) of TPR and (1-FPR). All metrics are reported as the mean values of 100 simulation replications, with bars indicating their standard errors.

**Figure 3:**
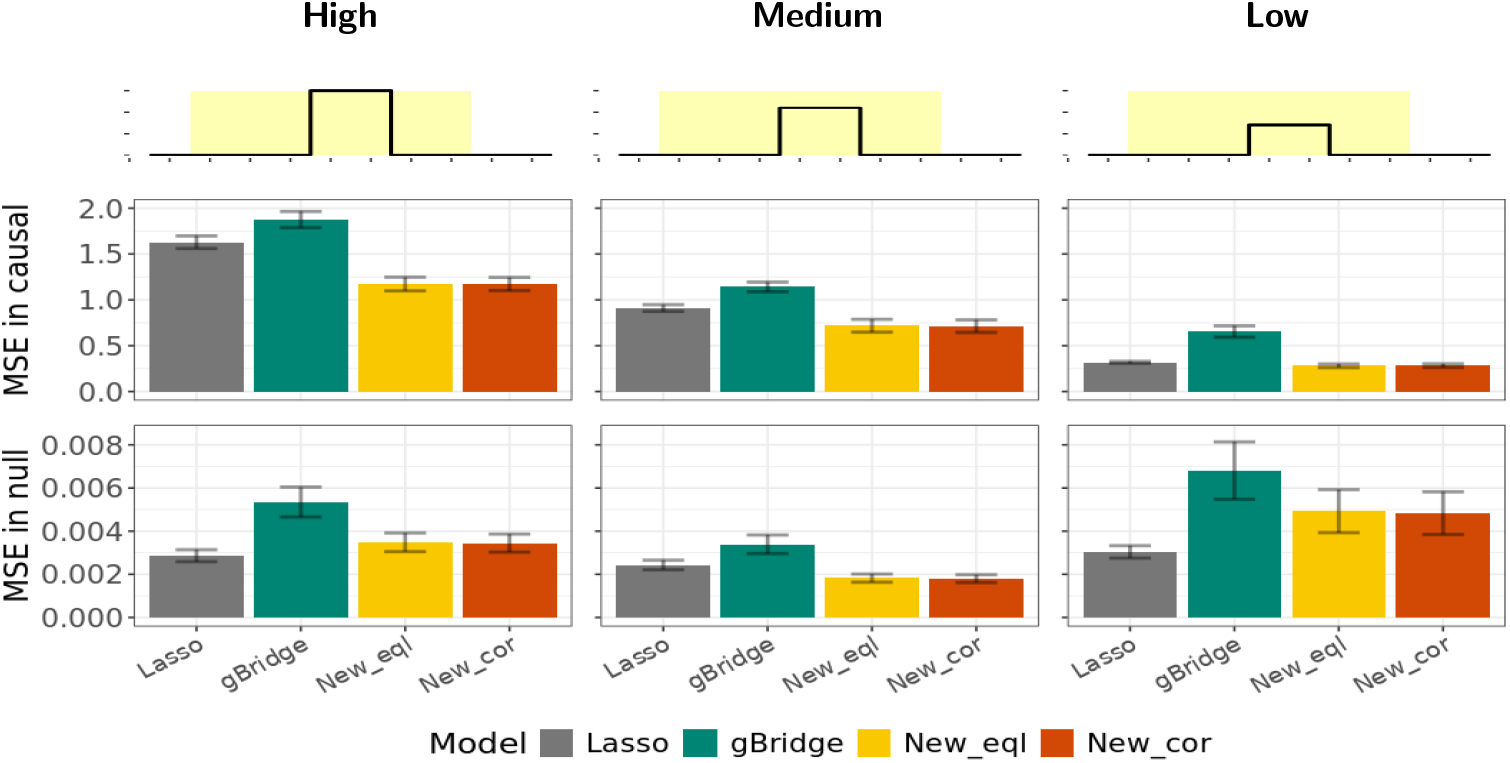
Effect estimation performance evaluated by mean squared error (MSE) under Simulation Scenario 1 for binary traits. The top row illustrates the design of causal signals under Scenario 1, with three levels of signal strength: high (left column), medium (middle column) to low (right column). The second and third rows show the MSEs in causal region and null (non-causal) regions, respectively. MSEs are reported as the mean values of 100 simulation replications, with bars indicating their standard errors.

Figure 4 (variable selection) and Figure 5 (effect estimation) show the results for binary traits under Scenario 2 (i.e., the entire region of consecutive CNV fragments being causal). The relative performance of different methods is similar to that observed under Scenario 1 but with two key differences. (a) The TPRs of `gBridge` are higher than `Lasso`, although they are still lower than the proposed methods. (b) Compared to `Lasso`, the FPRs of the proposed methods vary from lower to comparable to higher as signal strength decreases. Nevertheless, across various signal strengths, the proposed methods consistently yield better overall selection performance (i.e., higher G-means) and more accurate estimates of causal effects (i.e., lower MSEs) than `Lasso` and `gBridge`.

**Figure 4:**
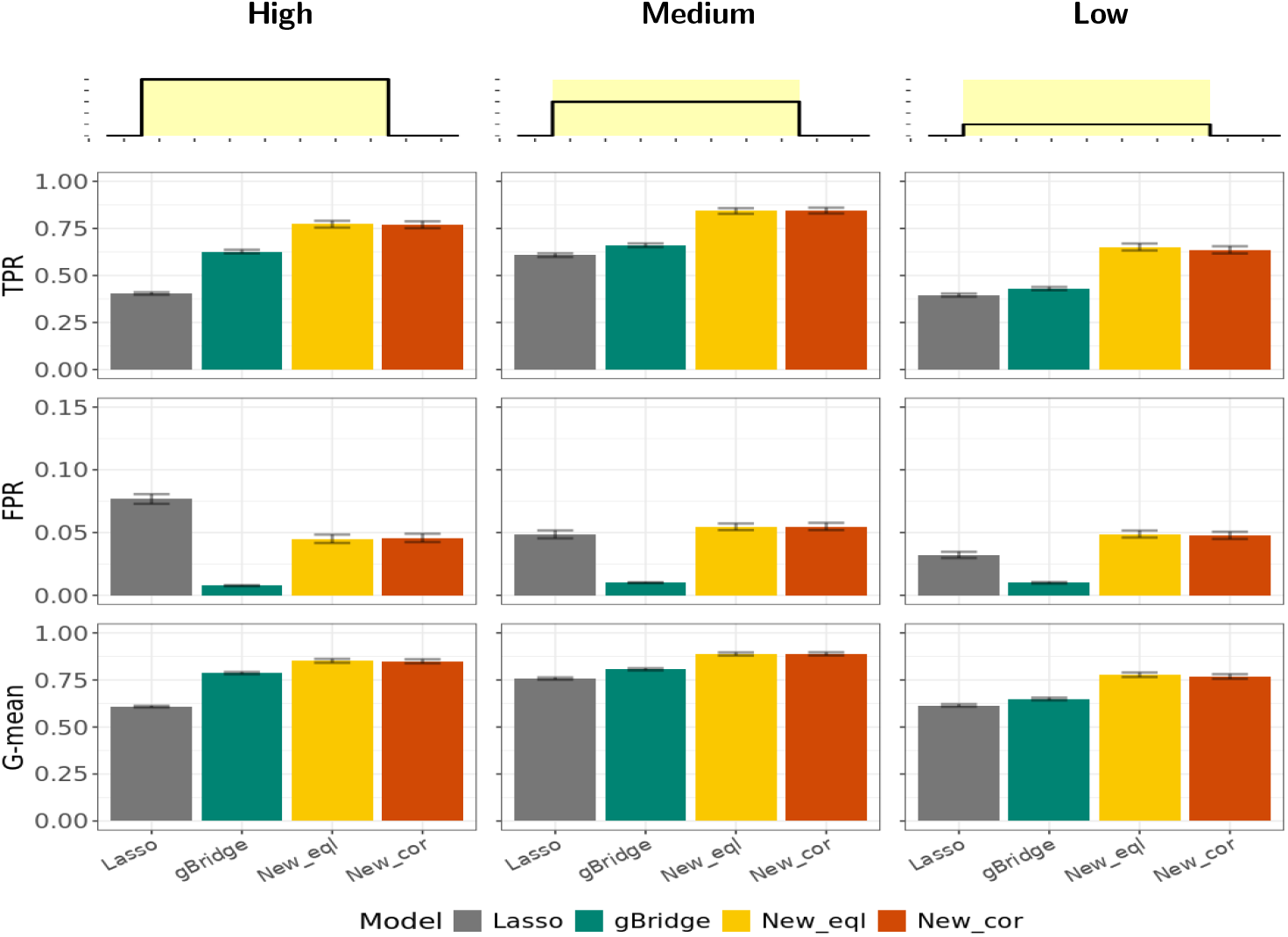
Variable selection performance evaluated by TPR, FPR, and G-mean under Simulation Scenario 2 for binary traits. The top row illustrates the design of causal signals under Scenario 2, with three levels of signal strength: high (left column), medium (middle column) to low (right column). The second to fourth rows respectively show the true positive rate (TPR), the false positive rate (FPR), and the geometric mean (G-mean) of TPR and (1-FPR). All metrics are reported as the mean values of 100 simulation replications, with bars indicating their standard errors.

**Figure 5:**
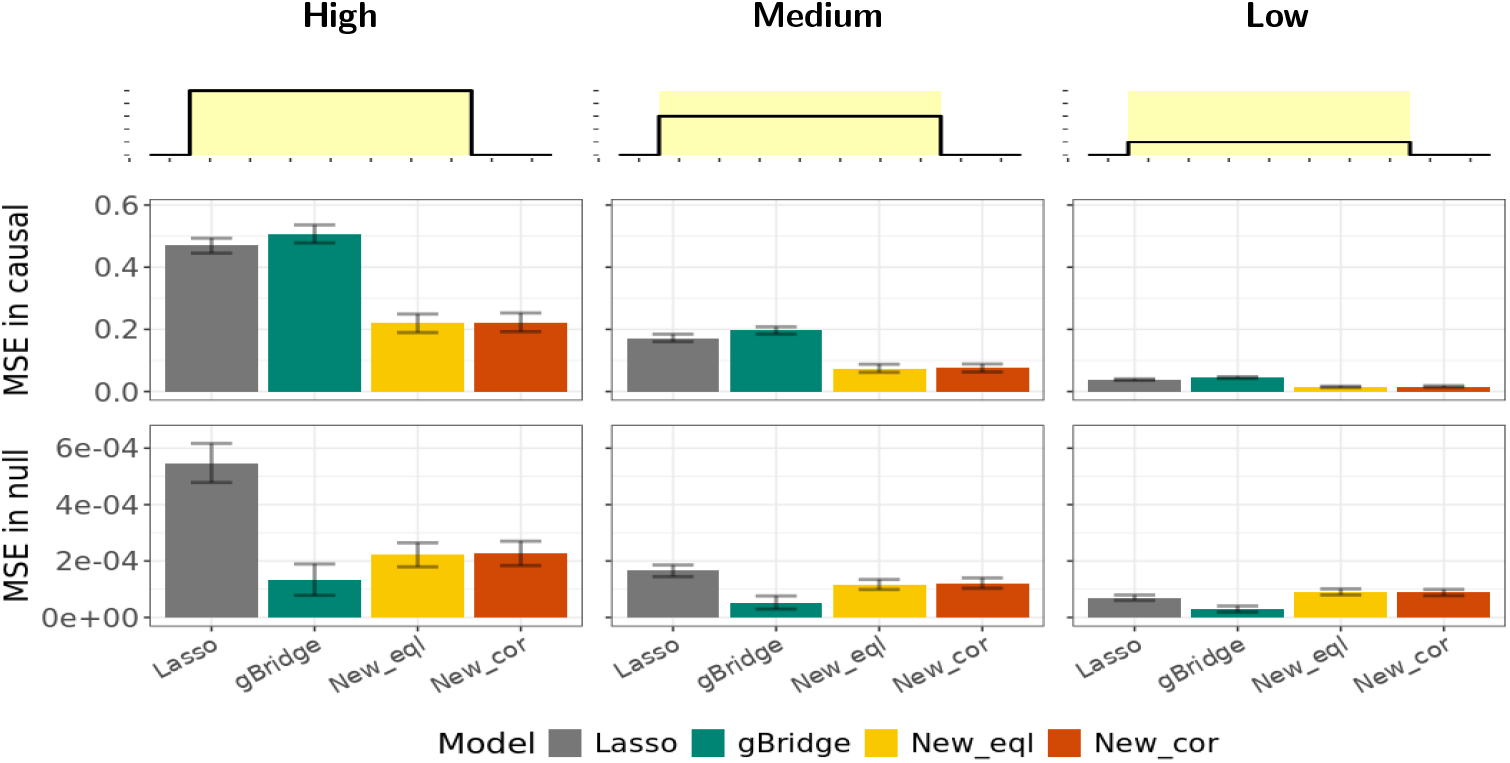
Effect estimation performance evaluated by mean squared error (MSE) under Simulation Scenario 2 for binary traits. The top row illustrates the design of causal signals under Scenario 2, with three levels of signal strength: high (left column), medium (middle column) to low (right column). The second and third rows show the MSEs in causal region and null (non-causal) regions, respectively. MSEs are reported as the mean values of 100 simulation replications, with bars indicating their standard errors.

Figure 6 (variable selection) and Figure 7 (effect estimation) show the results for binary traits under Scenario 3 (i.e., two sub-regions of consecutive CNV fragments being causal and with different effect sizes). The relative performance of different methods is similar to that observed in Scenario 2, except that the differences in variable selection performance among different methods are much smaller. Scenario 3 is more challenging for the proposed methods because the consecutive pattern of causal signals is interrupted. Nevertheless, the proposed methods still yield comparable overall selection performance and better effect estimates compared to the baseline methods, highlighting the adaptivity of the smooth penalty to the underlying true effect mechanisms.

**Figure 6:**
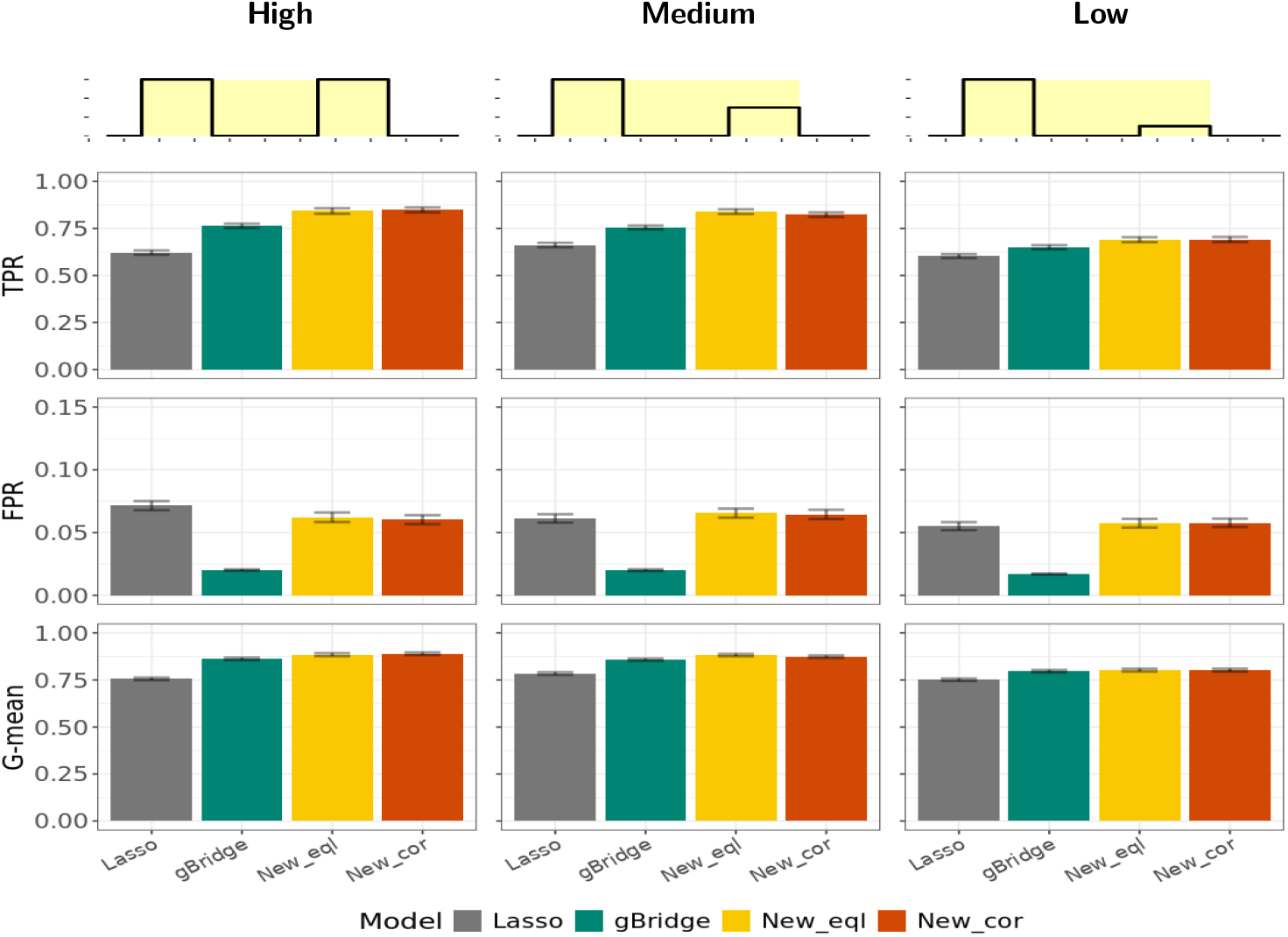
Variable selection performance evaluated by TPR, FPR, and G-mean under Simulation Scenario 3 for binary traits. The top row illustrates the design of causal signals under Scenario 3, with three levels of signal strength: high (left column), medium (middle column) to low (right column). The second to fourth rows respectively show the true positive rate (TPR), the false positive rate (FPR), and the geometric mean (G-mean) of TPR and (1-FPR). All metrics are reported as the mean values of 100 simulation replications, with bars indicating their standard errors.

**Figure 7:**
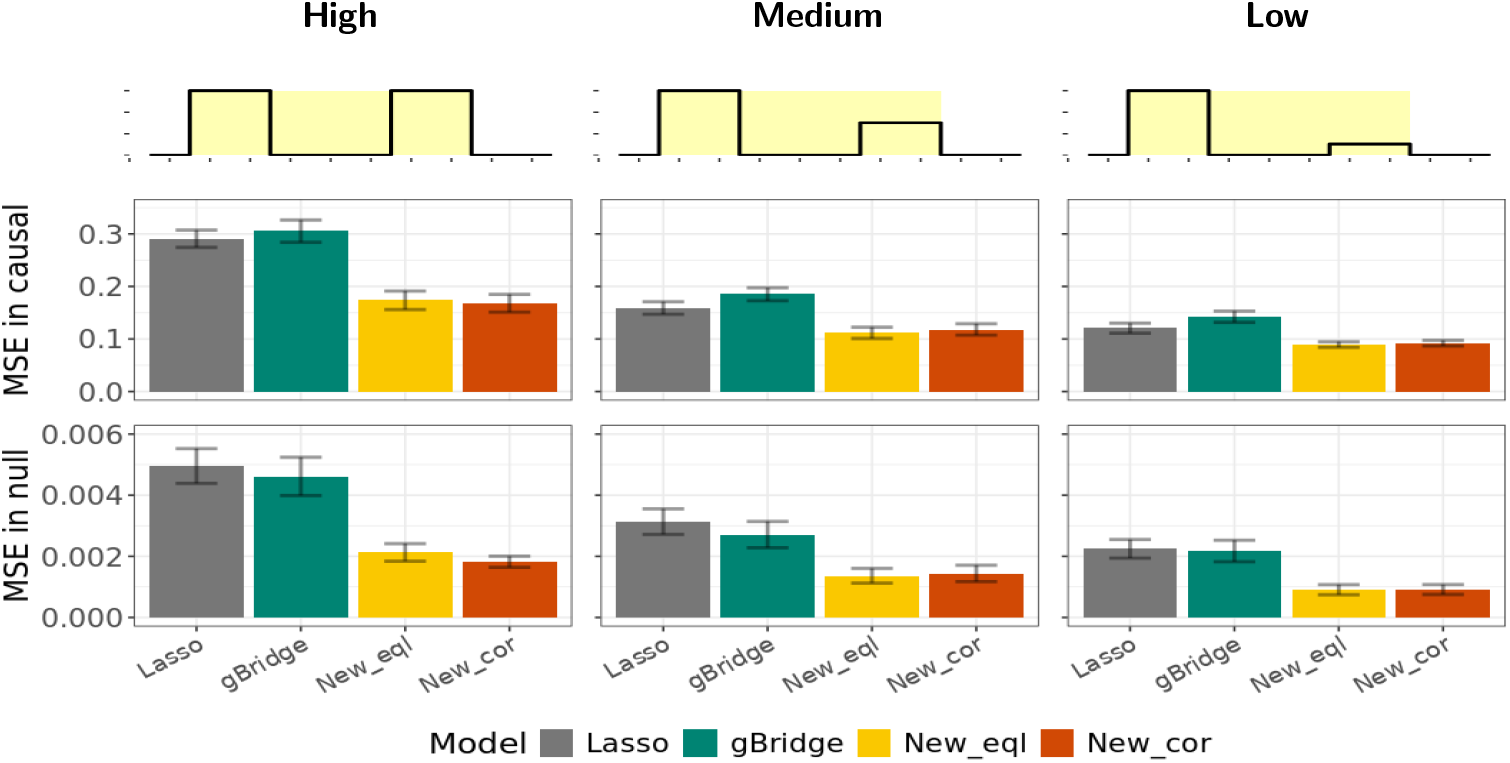
Effect estimation performance evaluated by mean squared error (MSE) under Simulation Scenario 3 for binary traits. The top row illustrates the design of causal signals under Scenario 3, with three levels of signal strength: high (left column), medium (middle column) to low (right column). The second and third rows show the MSEs in causal region and null (non-causal) regions, respectively. MSEs are reported as the mean values of 100 simulation replications, with bars indicating their standard errors.

The results of the continuous traits are shown in the supplemental information (**????????????**), and the findings are consistent with binary traits. In summary, the proposed models can select more true signals while keeping FPR at reasonable levels, result in a better or comparable overall selection performance, and yield more accurate effect size estimates.

### Real data application

Figure 8 shows the AD-associated CNV fragments identified by different methods, where “set size” indicates the number of associated CNV fragments identified by each method.

**Figure 8:**
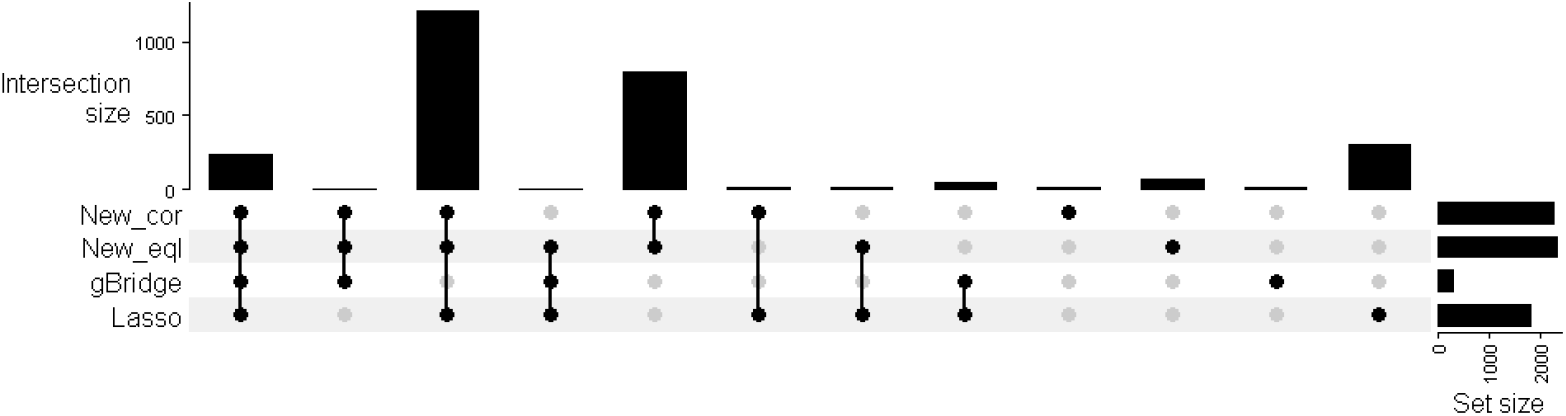
Number of AD-associated CNV fragments selected by different methods.

The proposed methods (i.e., `New_eql` and `New_cor`) select the largest number of associated CNVs (2,515 and 2,440 fragments, respectively), followed by `Lasso` (1,905) and then `gBridge` (299). For the proposed methods, the two different weighting schemes of effect smoothing selected a comparable number of CNV fragments. These results are consistent with those from the simulation studies.

Compared the genes overlapping the identified CNVs from each method with the known AD risk genes reported by the ADSP Gene Verification Committee, all methods identify AD-associated CNVs near *SPPL2A*. `Lasso`, `New_eql` and `New_cor` also identify associated CNVs near *NCK2, MME, JAZF1* and *RBCK1*. The proposed methods, `New_eql` and `New_cor`, additionally identify associated CNVs near *INPP5D, MINDY2*, and *ADAM10*. All of the above-mentioned AD-associated genes are supported by genomewide significance evidence in the literature^49–51^. Moreover, there is emerging evidence to show the potential roles of these genes in AD pathology, such as in the metabolism and accumulation of A*β* (e.g., *SPPL2A*^52^, *MME* ^53^, *ADAM10* ^50^, and *INPP5D* ^50^), tau phosphorylation (e.g., *JAZF1* ^54^), and neuron development and the regulation of synaptic transmission (e.g., *NCK2* ^55^). Additionally, AD-associated deletions near *NCK2* and *JAZF1* have also been reported in a recent study of AD-associated CNV calls from ADSP WGS data.^32^

Figure 9 shows the enriched biological processes in the GO enrichment analysis that have the Bonferroni adjusted p-values *<*0.05. `Lasso`, `New_eql` and `New_cor` identify 2 enriched biological processes in common (processes 1 and 2). `New_eql` additionally identifies processes 4 and 5 and `New_cor` additionally identifies processes 4-6. While not identifying any processes in common with other methods, `gBridge` identifies process 3.

**Figure 9:**
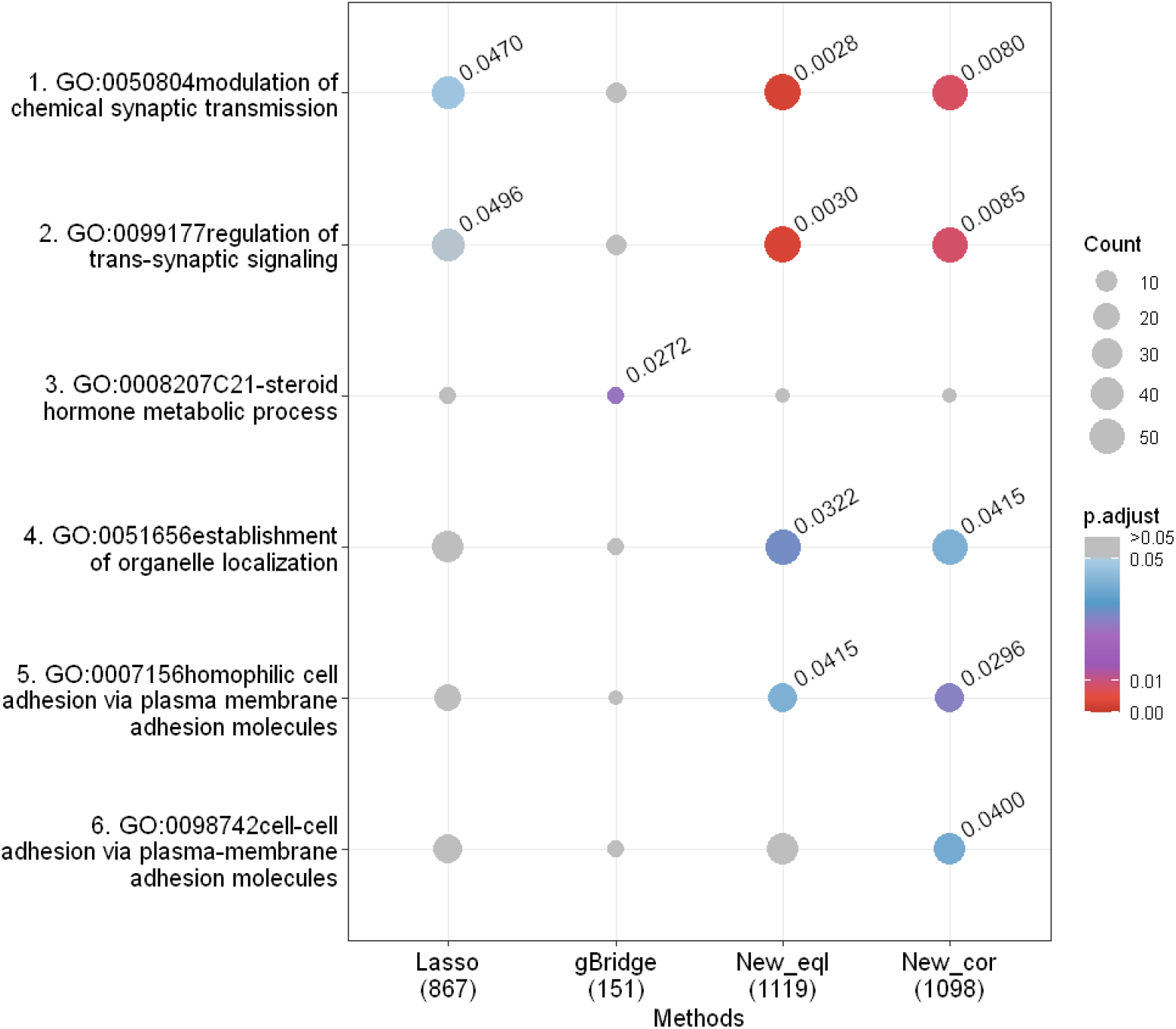
Gene ontology (GO) enrichment analysis for AD-associated genes identified by different methods. The dot color indicates the Bonferroni-adjusted p-value (p.adjust), and for significant GO terms, the adjusted p-values are shown next to the dots. The dot size indicates the count of genes identified by a certain method (X-axis) overlapping with each biological process (Y-axis). The number in parentheses on the X-axis indicates the number of genes with GO annotations for each method.

Biological processes 1 and 2 (commonly identified by `Lasso` and the proposed methods) are essential for neuron communication, involving intricate signaling mechanisms like neurotransmitter release and receptor activation.^56^. In AD, amyloid-*β* (A*β*) and tau serve as toxins in developing synaptic degeneration and impair neurotransmitter release and receptor-mediated activities, which consequently lead to memory and cognitive deficits^57,58^.

Process 3 (uniquely identified by `gBridge`) is related to the biosynthesis and metabolic of hormone molecules (e.g., aldosterone and progesterone)^59^; these hormones may play an essential role in AD pathogenesis and provides promising targets for AD management.^60,61^

Processes 4-6 (i.e., processes uniquely identified by the proposed methods) highlight key cellular mechanisms relevant to AD. Subcellular organelles such as endoplasmic reticulum, mitochondria, and lysosomes (process 4) are essential for neuron survival and synaptic function, with dysfunction leading to impaired energy production, protein misfolding, and disrupted calcium homeostasis, all contributing to AD pathology.^62–64^ Processes 5-6 focus on cell-cell adhesion in the nervous system. Dysregulation of adhesion molecules affects synaptic plasticity, A*β* metabolism, and neuroinflammation, contributing to AD progression.^65^

For the false negative assessment using “null” analysis, where we permute the AD status and generate 50 null datasets, the FPR for `Lasso` is 0.0059 with standard error (SE) 0.00074. For `gBridge`, `New_eql`, and `New_cor`, their FPRs (and SEs) are 0.0010 (0.00028), 0.0027 (0.00079), and 0.0027 (0.00079), respectively. That is, the two proposed methods have the same, reasonable FPRs, and as observed in the simulations, their FPRs are larger than `gBridge` but smaller than `Lasso`.

## Discussion

We propose a CNV profile regression framework to evaluate CNV association. The proposed method adopts a joint modeling framework to assess the effects of all CNVs over a genomic region (e.g., chromosome) simultaneously, uses the Lasso penalty to select associated CNVs and a weighted fusion penalty to encourage CNVs of adjacent positions to have similar effects on traits. We provide an R package “*CNVreg* “ for implementing the proposed methods, which accepts PLINK CNV format as input and outputs the CNV effect at each unit-length position (e.g., bp or 100bps).

The underlying rationale of CNV profile regression is to model an individual’s CNV events as a piecewise constant curve (i.e., CNV profile curve^27^), and the corresponding CNV effects can be summarized as a “CNV effect curve”. There are several attractive features for assessing CNV association using an individual’s entire CNV profile. (a) The CNV profile curve naturally captures the CNV information in dosage and length. (b) It bypasses the need to define CNV loci when evaluating association. (c) Under the assumption of constant effect within a CNV fragment^19^, the CNV profile regression can be expressed as the typically considered fragment-based regression, except that the predictor 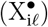 is dosage×length, i.e., the area under the CNV profile curve for duplications/deletions within each fragment. (d) Using the data augmentation approaches^66–68^ as described in Appendix A, the proposed method can be effectively implemented using standard packages for penalized regression such as *glmnet* ^34^. (e) Compared to typical fragment-based association models, where the predictor is the dosage (instead of dosate×length) in a fragment and the results depend on study-specific fragment definitions, the effect sizes from the proposed method are directly comparable across different studies regardless of fragment definitions. This is because the proposed method reports the effect size at each unit-length position, although adjacent positions can have identical effect sizes if they are within the same fragment. (f) The proposed method also illustrates how to convert typical fragment-based results (e.g., *α*_*ℓ*_’s in Equation (5)) for cross-study comparisons. The key is to divide the “aggregate” effect size (*α*_*ℓ*_) by the length of the CNV fragment and obtain the effect size per unit length. Such length-adjusted effect can then be used for direct comparisons or meta-analysis across different studies.

Numerical evaluations using simulated and real data show that the proposed methods (`New_eql` and `New_cor`) have better or comparable performance in variable selection and better effect size estimates than the baseline methods. Compared to `Lasso`, we see that using weighted fusion penalty to account for adjacent CNV structures can increase TPRs while maintaining similar or lower FPRs and reduce estimation errors. In our analysis, both equal weight and correlation weight often produce similar results, suggesting that the adjacent CNV fragments are often correlated at similar levels and hence correlation weights are nearly equal. Although `gBridge` also incorporates adjacent CNV structures, it gives more conservative results, e.g., selecting a smaller number of CNVs, resulting in the lowest (best) FPRs but meanwhile lower TPRs and higher MSEs for causal effect estimates.

The proposed methods have some limitations. First, fitting CNV profile regression requires two-dimensional grid search for selecting the optimal tuning parameters (λ_1_, λ_2_), and hence the computational cost is higher than the baseline methods. This cost can be substantial for extremely large WGS datasets. To address this issue, our R package provides options for parallel computing, allowing users to distribute tasks across multiple CPU nodes. Specifically, users can perform different folds of cross-validation and fit regression models for different candidate pairs of (λ_1_, λ_2_) in parallel, thereby reducing runtime and improving computational efficiency. Second, the proposed framework does not provide p-values. It is possible to conduct post-selection inference procedures to obtain p-values if needed. One approach is to randomly split the data into two equal-sized subsets: one subset for conducting penalized CNV profile regression, and the other for fitting the model obtained from variable selection using traditional linear or logistic regression methods. This approach allows the use of standard inference techniques to assess the significance of selected CNVs, although it comes at the cost of reducing the effective sample size by half.

## Supporting information

All supplemental

## APPENDIX A Fitting penalized CNV profile regression for continuous and binary traits

To estimate the model parameters, we adapt the data augmentation approach as in elastic net^66,67^ and fused Lasso^68^. That is, we rewrite Equation (6) into the following form using carefully defined response variable Y^∗^ and predictors X^∗^ and Z^∗^:

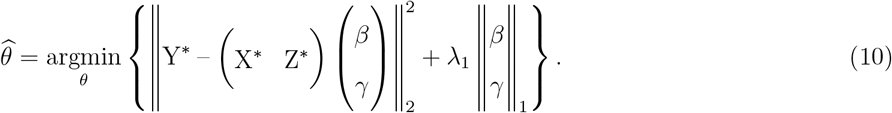

Consequently Equation (10) can be solved as a Lasso regression using existing software such as R package *glmnet*.

For continuous traits (CT), we set 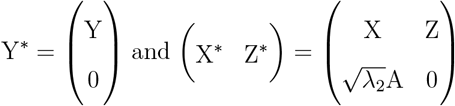. Then *θ* can be obtained by solving

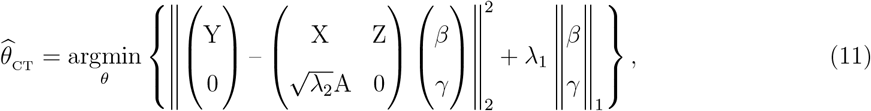

For binary traits (BT), we iteratively perform least squares to solve penalized logistic regression. Let 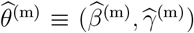 be the estimates of *θ* = (*β, γ*) at the m-th iteration. The quadratic approximation of the log-likelihood function, using the Taylor’s expansion at 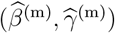,is indeed a penalized weighted least square^67,68^. Specifically, define

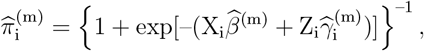

the working response 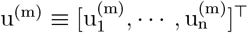 where

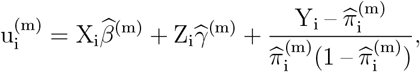

and the weight matrix Ω^(m)^

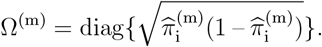

Then in the (m + 1)th iteration, we set 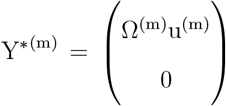 and 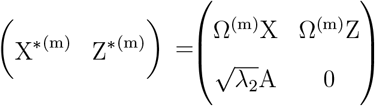. We can update *θ* by solving

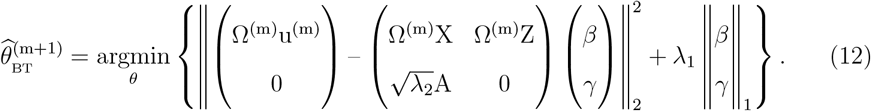

## Web resources

1. The Ensembl Variant Effect Predictor (VEP) website: https://useast.ensembl.org/Homo_sapiens/Tools/VEP?db=core
2. The QuickGo website: https://www.ebi.ac.uk/QuickGO/
3. List of AD loci and genes with genetic evidence compiled by ADSP Gene Verification Committee: https://adsp.niagads.org/gvc-top-hits-list/
4. Github link to R package ‘*CNVreg* ‘: https://github.com/Oceanyq/CNVreg

## Data and code availability

Simulation Code:

https://github.com/Oceanyq/CNVreg/tree/main/Simulation%20Code

## Author contributions

The authors’ contributions to the paper are as follows: study conceptualization and methodology: YS, WL, AB, JYT; data curation: YS, HW, AAT, AB, YC, LSW, GS, WPL, JYT; investigation, analysis, and visualization: YS, WL, SH, HW, AAT, YC, LSW, GS, WPL, JYT; software: YS, SH, YC, AAT, JYT; funding acquisition: WPL, JYT; writing – original draft: YS, WL, JYT; writing – review & editing: all authors reviewed, critically revised, and approved the manuscript.

## Acknowledgements

This work was supported by the U.S. National Institutes of Health RF1AG074328.

---

**Figure S1:**
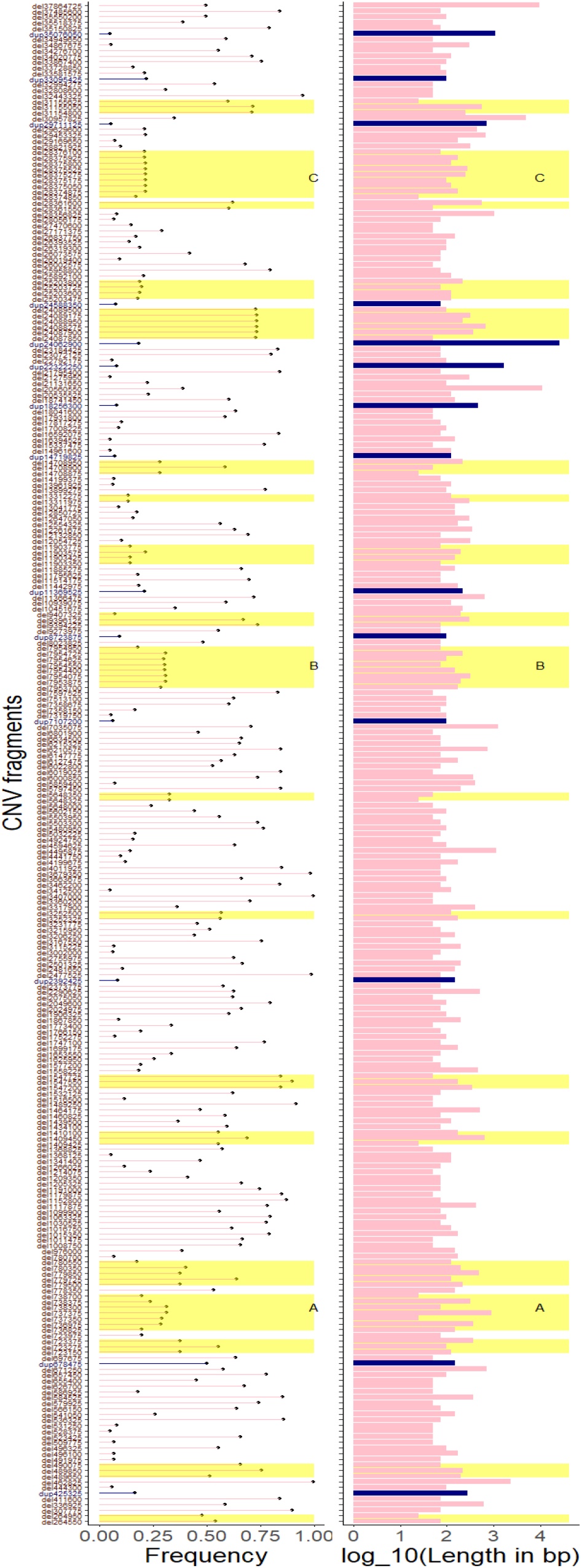
Frequency (left column) and length (right column) of CNV fragments used in simulation studies. The data is from Chromosome 10p of Non-Hispanic White (NHW) individuals in the ADSP dataset. In both columns, the y-axis indicates the CNV fragments; red indicates CNV deletion fragments; and blue indicates CNV duplication fragments. Yellow blocks highlight regions with consecutive CNV fragments. Blocks A, B, and C denote the candidate causal regions for setting causal signals in the simulation studies.

**Figure S2:**
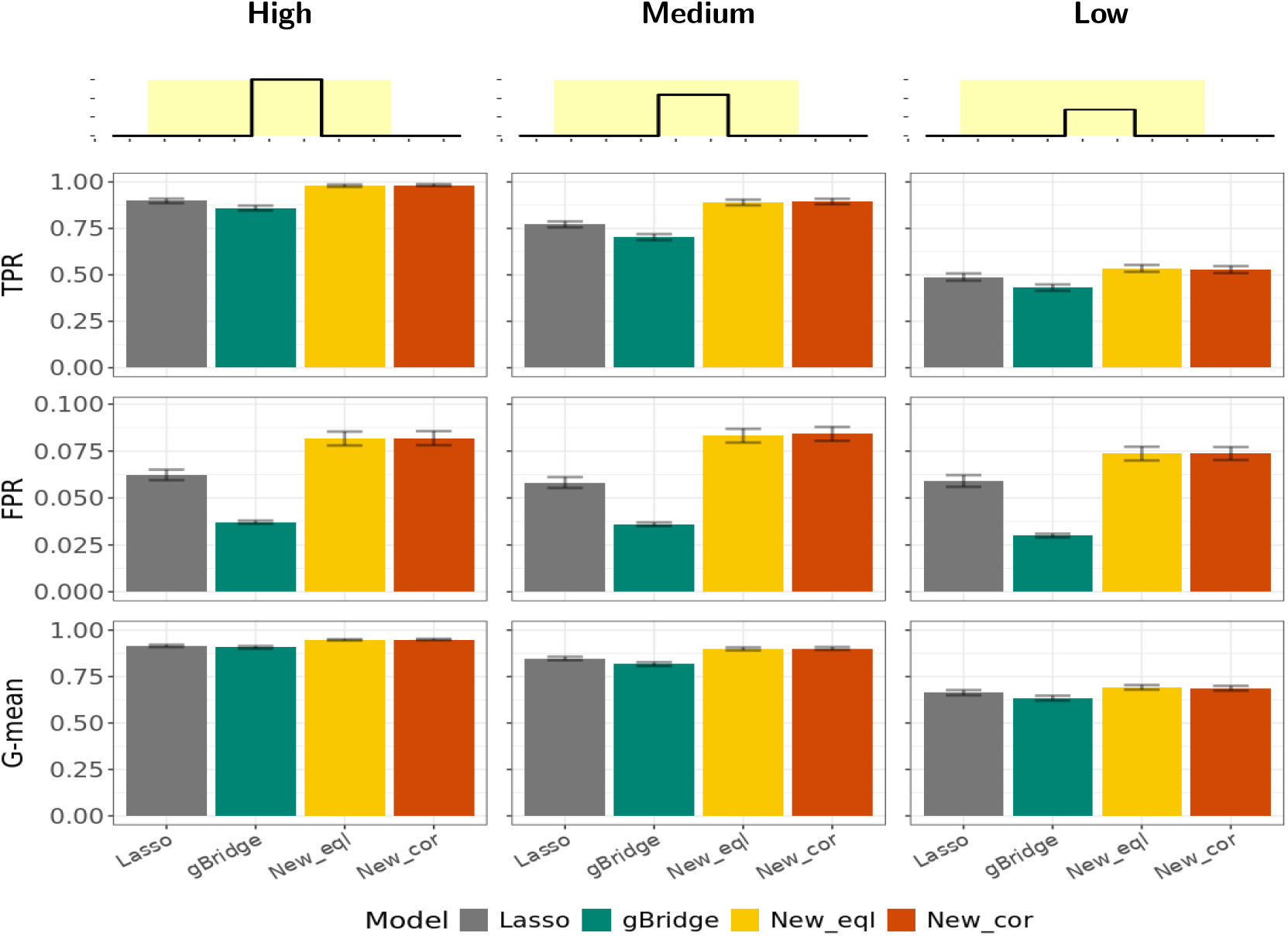
Variable selection performance evaluated by TPR, FPR, and G-mean under Simulation Scenario 1 for continuous traits. The top row illustrates the design of causal signals under Scenario 1, with three levels of signal strength: high (left column), medium (middle column) to low (right column). The second to fourth rows respectively show the true positive rate (TPR), the false positive rate (FPR), and the geometric mean (G-mean) of TPR and (1-FPR). All metrics are reported as the mean values of 100 simulation replications, with bars indicating their standard errors.

**Figure S3:**
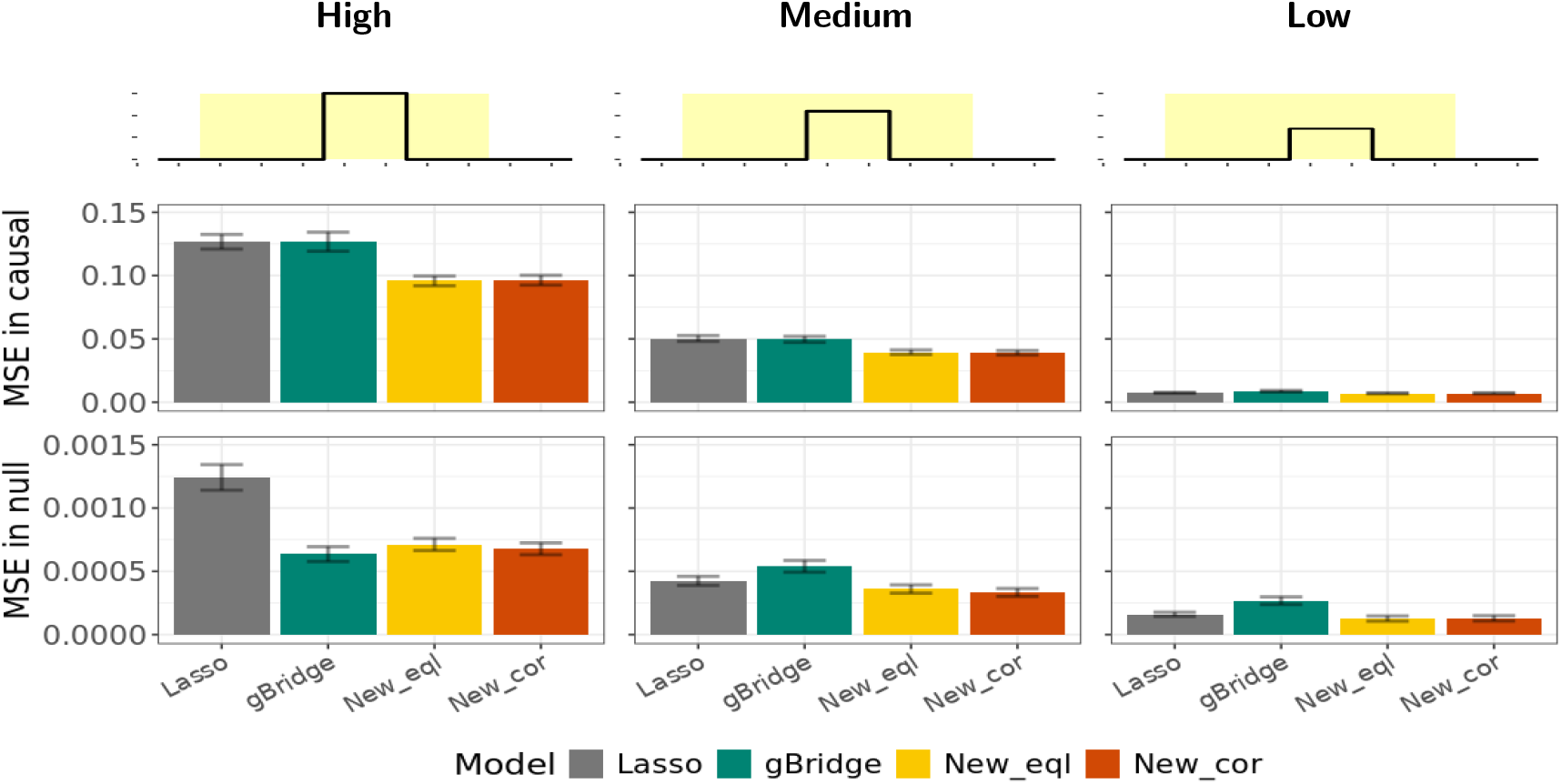
Effect estimation performance evaluated by mean squared error (MSE) under Simulation Scenario 1 for continuous traits. The top row illustrates the design of causal signals under Scenario 1, with three levels of signal strength: high (left column), medium (middle column) to low (right column). The second and third rows show the MSEs in causal region and null (non-causal) regions, respectively. MSEs are reported as the mean values of 100 simulation replications, with bars indicating their standard errors.

**Figure S4:**
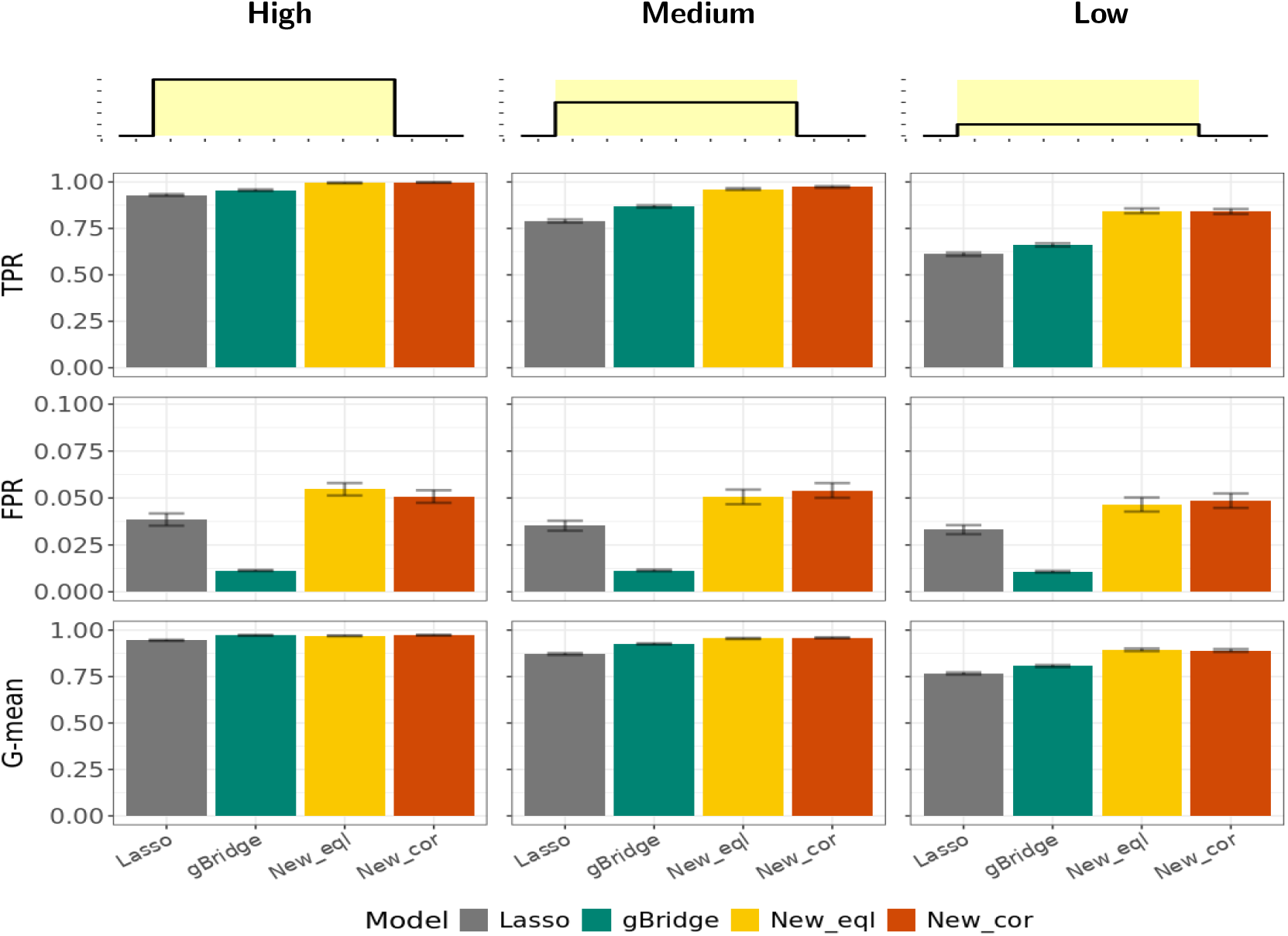
Variable selection performance evaluated by TPR, FPR, and G-mean under Simulation Scenario 2 for continuous traits. The top row illustrates the design of causal signals under Scenario 2, with three levels of signal strength: high (left column), medium (middle column) to low (right column). The second to fourth rows respectively show the true positive rate (TPR), the false positive rate (FPR), and the geometric mean (G-mean) of TPR and (1-FPR). All metrics are reported as the mean values of 100 simulation replications, with bars indicating their standard errors.

**Figure S5:**
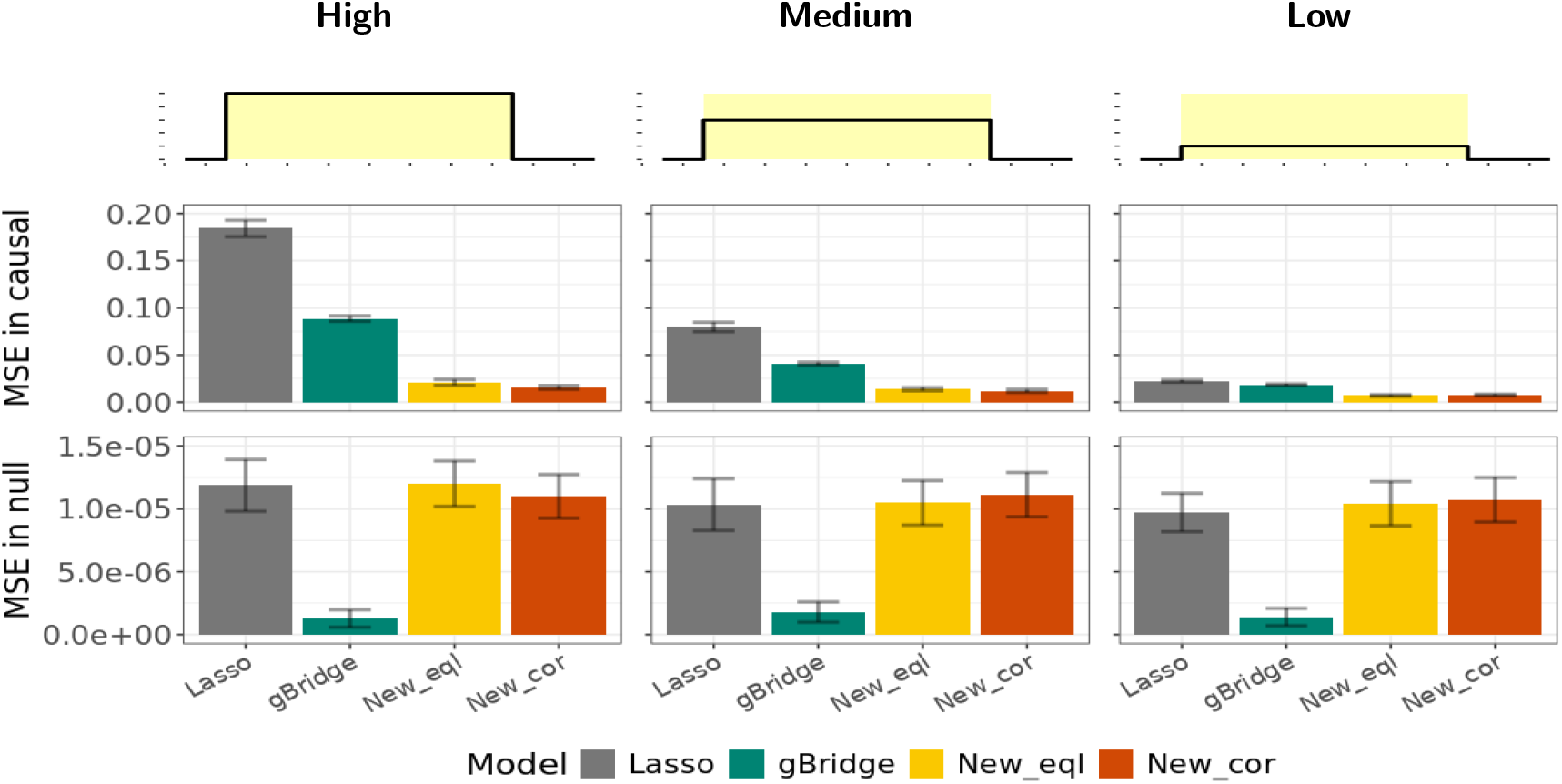
Effect estimation performance evaluated by mean squared error (MSE) under Simulation Scenario 2 for continuous traits. The top row illustrates the design of causal signals under Scenario 2, with three levels of signal strength: high (left column), medium (middle column) to low (right column). The second and third rows show the MSEs in causal region and null (non-causal) regions, respectively. MSEs are reported as the mean values of 100 simulation replications, with bars indicating their standard errors.

**Figure S6:**
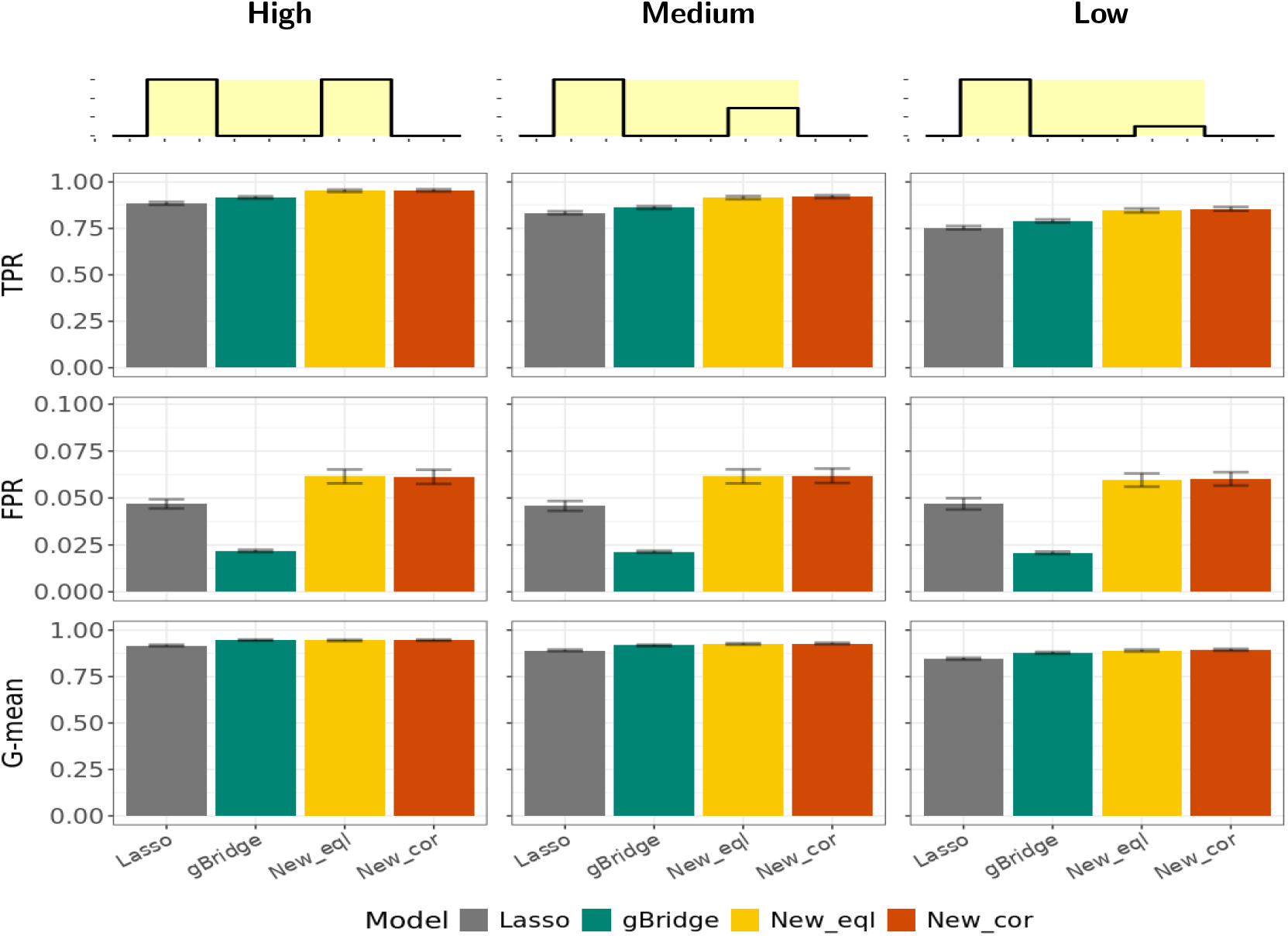
Variable selection performance evaluated by TPR, FPR, and G-mean under Simulation Scenario 3 for continuous traits. The top row illustrates the design of causal signals under Scenario 3, with three levels of signal strength: high (left column), medium (middle column) to low (right column). The second to fourth rows respectively show the true positive rate (TPR), the false positive rate (FPR), and the geometric mean (G-mean) of TPR and (1-FPR). All metrics are reported as the mean values of 100 simulation replications, with bars indicating their standard errors.

**Figure S7:**
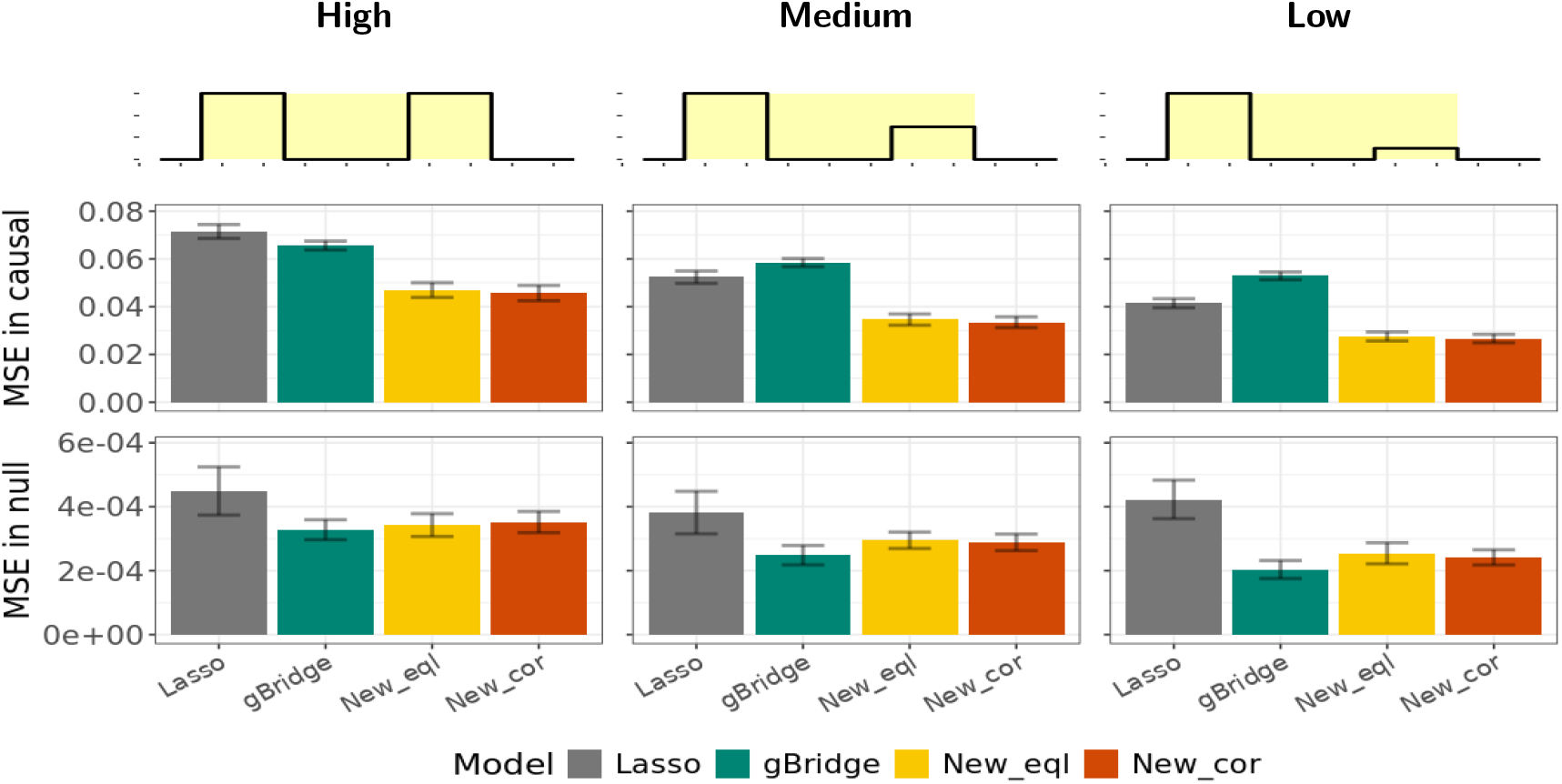
Effect estimation performance evaluated by mean squared error (MSE) under Simulation Scenario 3 for continuous traits. The top row illustrates the design of causal signals under Scenario 3, with three levels of signal strength: high (left column), medium (middle column) to low (right column). The second and third rows show the MSEs in causal region and null (non-causal) regions, respectively. MSEs are reported as the mean values of 100 simulation replications, with bars indicating their standard errors.

