## Supplementary material for "CNV-Profile Regression: A New Approach for Copy Number Variant Association Analysis in Whole Genome Sequencing Data": All supplemental

Si et al.

### Pseudo code for fitting penalized CNV profile regression with Lasso and weighted fusion penalties

**Algorithm 1:** $\text{cv\_fit}(X, Z, Y, A, M_{\lambda_1}, M_{\lambda_2}, K = 5)$

/* Use K-fold CV to select $\lambda_1, \lambda_2$ that minimizes the average validation loss, and return the final model at the selected $\lambda_1$ and $\lambda_2$. */

**Input** : CNV X; covariate Z; trait Y; weight matrix of CNV adjacency structure A; list of candidate $\lambda_1$'s $M_{\lambda_1}$; list of candidate $\lambda_2$'s $M_{\lambda_2}$; fold number in CV K

**Output:** The selected pair of $(\widehat{\lambda}_1, \widehat{\lambda}_2)$
Coefficient estimate $\widehat{\theta}$ at $(\widehat{\lambda}_1, \widehat{\lambda}_2)$

/* Randomly split data into K folds. */
$X = \{X_1, \ldots, X_K\}$;
$Z = \{Z_1, \ldots, Z_K\}$;
$Y = \{Y_1, \ldots, Y_K\}$;
/* For each pair of candidate $\lambda_1$ and $\lambda_2$, iteratively use 1 of the K folds as validation set and the other folds as training set */
**for** $\lambda_1 \in M_{\lambda_1}$ **do**
  **for** $\lambda_2 \in M_{\lambda_2}$ **do**
    **for** $i = 1$ **to** K **do**
      /* Split training data and validation data */
      $X_{tr} = X \backslash \{X_i\}$, $X_{val} = X_i$;
      $Z_{tr} = Z \backslash \{Z_i\}$, $Z_{val} = Z_i$;
      $Y_{tr} = Y \backslash \{Y_i\}$, $Y_{val} = Y_i$;
      **if** *Y is a continuous trait* **then**
        /* Fit the penalized regression for continuous trait */
        $\widehat{\theta} = \text{CT\_lasso}(X_{tr}, Z_{tr}, Y_{tr}, A, \lambda_1, \lambda_2)$ (See Algorithm 2);
      **else if** *Y is a binary trait* **then**
        /* Fit the penalized regression for binary trait */
        $\widehat{\theta} = \text{BT\_lasso}(X_{tr}, Z_{tr}, Y_{tr}, A, \lambda_1, \lambda_2)$ (See Algorithm 3);
      **end if**
      /* Calculate loss in validation set */
      $\text{loss}_i^{\lambda_1,\lambda_2} = \text{Loss}([X_{val} Z_{val}], Y_{val}, \widehat{\theta})$ (See Algorithm 4);
    **end for**
  **end for**
**end for**
/* Calculate average validation loss for all pairs of $\lambda_1$ and $\lambda_2$ */
$\text{loss}_{\text{Avg}}^{\lambda_1,\lambda_2} = \text{Average}(\text{loss}_i^{\lambda_1,\lambda_2}, i = [1, 2, 3, 4, 5])$;
/* Find the pair ($\lambda_1$, $\lambda_2$) that minimizes the average validation loss */
$(\widehat{\lambda}_1, \widehat{\lambda}_2)$ corresponds to $\lambda_1, \lambda_2$ that give the $\min(\text{loss}_{\text{Avg}}^{\lambda_1,\lambda_2})$;
/* Fit the final model at $(\widehat{\lambda}_1, \widehat{\lambda}_2)$ */
**if** *Y is a continuous trait* **then**
  $\widehat{\theta} = \text{CT\_lasso}(X, Z, Y, A, \widehat{\lambda}_1, \widehat{\lambda}_2)$;
**else if** *Y is a binary trait* **then**
  $\widehat{\theta} = \text{Bt\_las}(X, Z, Y, A, \widehat{\lambda}_1, \widehat{\lambda}_2)$;
**end if**
**return** $(\widehat{\lambda}_1, \widehat{\lambda}_2), \widehat{\theta}$

**Algorithm 2:** CT_lasso(X, Z, Y, A, $\lambda_1$, $\lambda_2$)

/* For a continuous trait Y, we directly apply Lasso regression to the augmented data ($Y^*$ and $X^*$) to estimate regression coefficients. */

**Input :** CNV X; covariate Z; trait Y; weight matrix of CNV adjacency structure A; candidate tuning parameters $\lambda_1, \lambda_2$

**Output:** Coefficient estimate $\widehat{\theta}$

/* Data augmentation */

$X^* = \begin{pmatrix} X & Z \\ \sqrt{\lambda_2}A & 0 \end{pmatrix}$;

$Y^* = \begin{pmatrix} Y \\ 0 \end{pmatrix}$;

/* Fit Lasso regression with the augmented data. Many well-established packages are available */

/* For example, use R package *glmnet* to fit lasso regression, and obtain the coefficient estimates at $\lambda_1$ */

fit $=$ glmnet($X^*, Y^*$);

$\widehat{\theta} = \text{coef}(\text{fit}, \lambda_1)$;

**return** $\widehat{\theta}$

**Algorithm 3:** BT_lasso$(\mathrm{X}, \mathrm{Z}, \mathrm{Y}, \mathrm{A}, \lambda_1, \lambda_2)$

`/* For a binary trait Y, we iteratively update the coefficient estimate` $\widehat{\theta}$ `with Lasso regression until converge. */`

**Input :** CNV X; covariate Z; trait Y; weight matrix of CNV adjacency structure A; candidate tuning parameters $\lambda_1, \lambda_2$

**Output:** Coefficient estimate $\widehat{\theta}$

`/* Initialize coefficient estimate` $\widehat{\theta}^{(0)}$`. */`

$\mathrm{m} = 0$;

$\mathrm{fit} = \mathrm{glmnet}([\mathrm{XZ}], \mathrm{Y}, \mathrm{family} = \text{“binomial”})$;

$\widehat{\theta}^{(0)} = \mathrm{coef}(\mathrm{fit}, \lambda_1)$;

`/* Initialize control parameters */`

*max_theta_diff* = inf;

*loss_diff* = inf;

`/* Iteratively update coefficient estimate until converge */`

**while** *max_theta_diff* $> 10^{-3}$ & *loss_diff* $> 10^{-6}$ **do**

- Obtain weight $\Omega^{(\mathrm{m})}$ and working response $\mathrm{u}^{(\mathrm{m})}$ as described in Appendix A;
- Construct weighted data $\mathrm{X}^{(\mathrm{m})} = \Omega^{(\mathrm{m})}\mathrm{X}, \mathrm{Z}^{(\mathrm{m})} = \Omega^{(\mathrm{m})}\mathrm{Z}$, and $\mathrm{Y}^{(\mathrm{m})} = \Omega^{(\mathrm{m})}\mathrm{u}^{(\mathrm{m})}$;
- `/* update coefficient estimate` $\widehat{\theta}^{(\mathrm{m}+1)}$ `*/`
- $\widehat{\theta}^{(\mathrm{m}+1)} = \mathrm{CT\_lasso}(\mathrm{X}^{(\mathrm{m})}, \mathrm{Z}^{(\mathrm{m})}, \mathrm{Y}^{(\mathrm{m})}, \mathrm{A}, \lambda_1, \lambda_2)$ (see Algorithm 2);
- $\mathrm{loss}^{(\mathrm{m}+1)} = \mathrm{Loss}([\mathrm{X}, \mathrm{Z}], \mathrm{Y}, \widehat{\theta}^{(\mathrm{m}+1)})$ (see Algorithm 4);
- *max_theta_diff* $= \max\left[|\widehat{\theta}^{(\mathrm{m}+1)} - \widehat{\theta}^{(\mathrm{m})}|\right]$;
- *loss_diff* $= |\mathrm{loss}^{(\mathrm{m}+1)} - \mathrm{loss}^{(\mathrm{m})}|$;
- $\widehat{\theta} = \theta^{(\mathrm{m}+1)}$;
- $\mathrm{m} = \mathrm{m} + 1$;

**end while**

**return** $\widehat{\theta}$

**Algorithm 4:** Loss$(\mathrm{X}, \mathrm{Z}, \mathrm{Y}, \widehat{\theta})$

`/* For continuous traits, use residual sum of squares as loss, and for binary traits, use negative log-likelihood as loss. */`

**Input :** CNV X; covariate Z; trait Y

**Output:** Estimated loss

**if** *Y is a continuous trait* **then**

- $\mathrm{loss} = \|\mathrm{y} - [\mathrm{XZ}]\widehat{\theta}\|_2^2$;

**else if** *Y is a binary trait* **then**

- $\mathrm{loss} = -\log(\mathrm{L}([\mathrm{XZ}] \equiv -\sum_{\mathrm{i}=1}^{\mathrm{n}} \{\mathrm{Y_i}([\mathrm{X_i Z_i}]\widehat{\theta}) - \log(1 + \exp([\mathrm{X_i Z_i}]\widehat{\theta}))\}$

**end if**

**return** loss

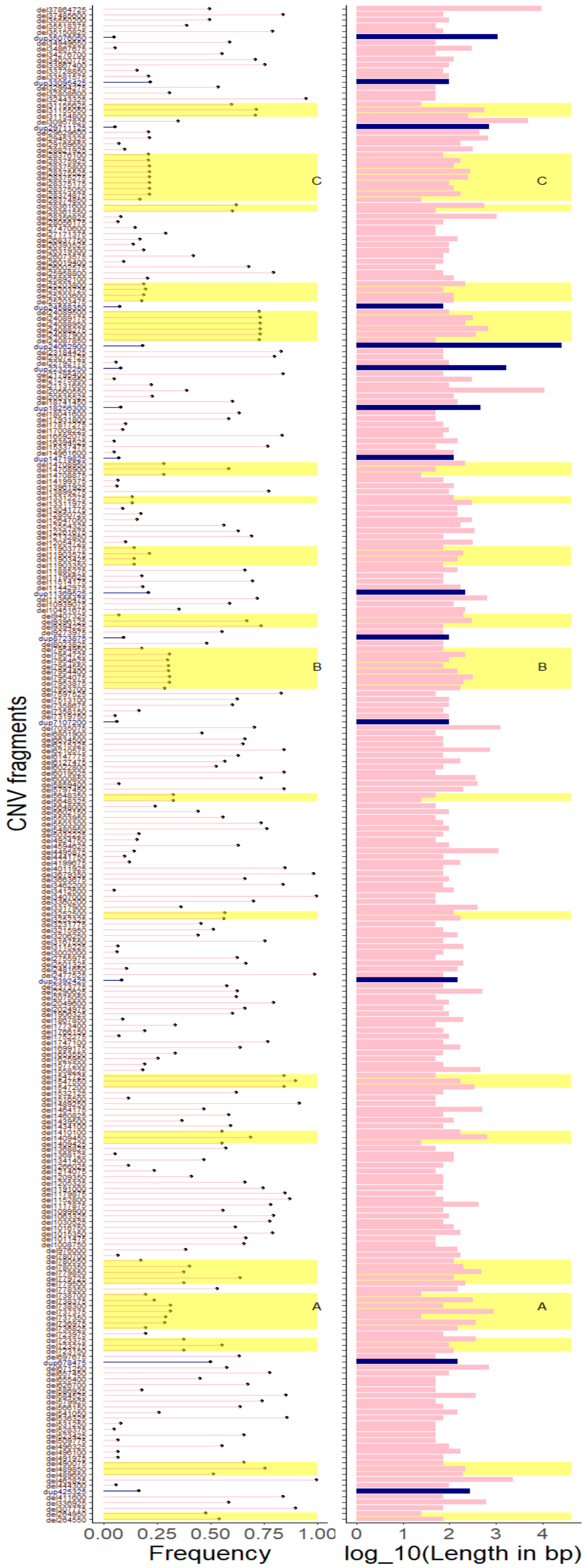


**Figure S1: Frequency (left column) and length (right column) of CNV fragments used in simulation studies.** The data is from Chromosome 10p of Non-Hispanic White (NHW) individuals in the ADSP dataset. In both columns, the y-axis indicates the CNV fragments; red indicates CNV deletion fragments; and blue indicates CNV duplication fragments. Yellow blocks highlight regions with consecutive CNV fragments. Blocks A, B, and C denote the candidate causal regions for setting causal signals in the simulation studies.

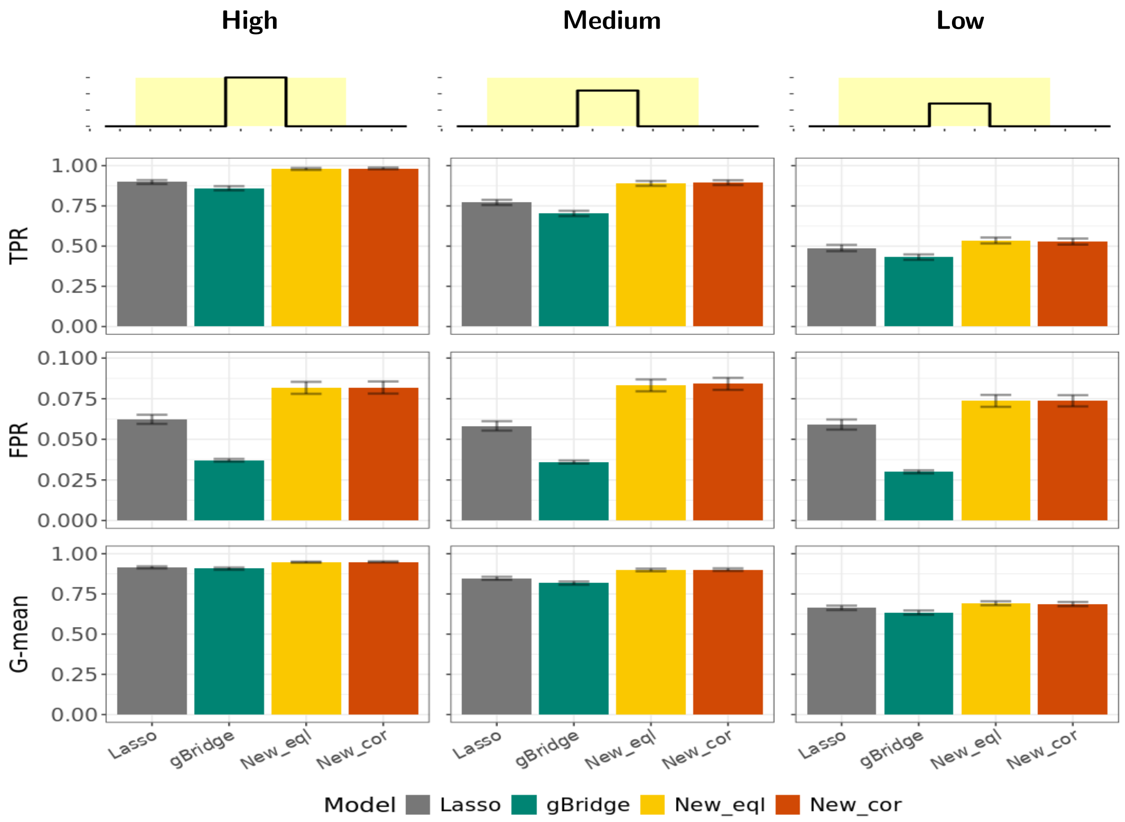


**Figure S2: Variable selection performance evaluated by TPR, FPR, and G-mean under Simulation Scenario 1 for continuous traits.** The top row illustrates the design of causal signals under Scenario 1, with three levels of signal strength: high (left column), medium (middle column) to low (right column). The second to fourth rows respectively show the true positive rate (TPR), the false positive rate (FPR), and the geometric mean (G-mean) of TPR and (1-FPR). All metrics are reported as the mean values of 100 simulation replications, with bars indicating their standard errors.

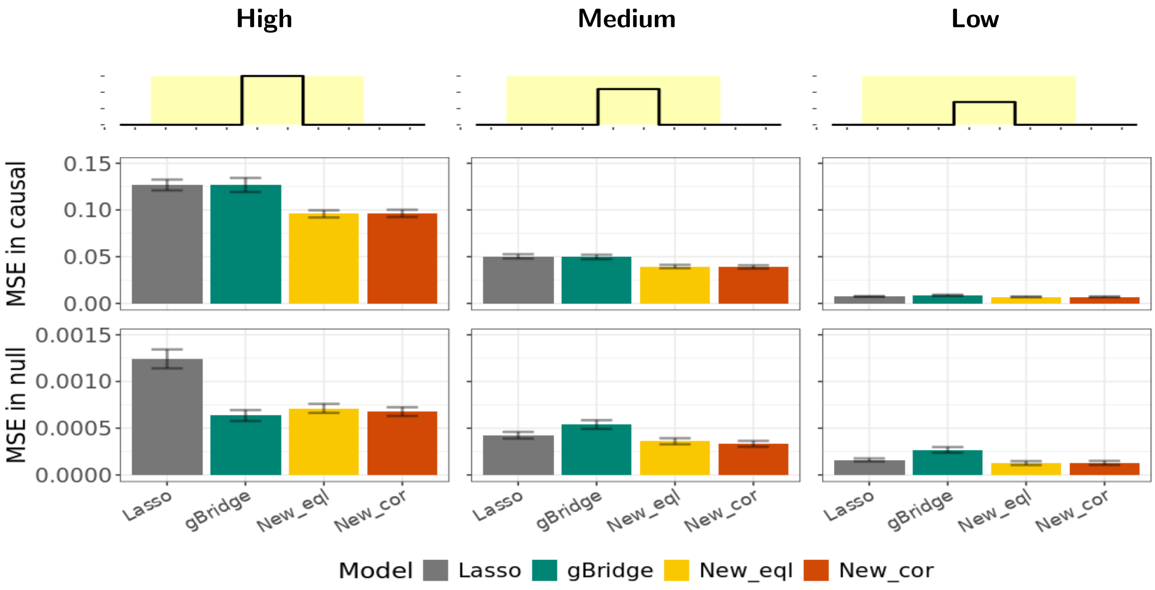


**Figure S3: Effect estimation performance evaluated by mean squared error (MSE) under Simulation Scenario 1 for continuous traits.** The top row illustrates the design of causal signals under Scenario 1, with three levels of signal strength: high (left column), medium (middle column) to low (right column). The second and third rows show the MSEs in causal region and null (non-causal) regions, respectively. MSEs are reported as the mean values of 100 simulation replications, with bars indicating their standard errors.

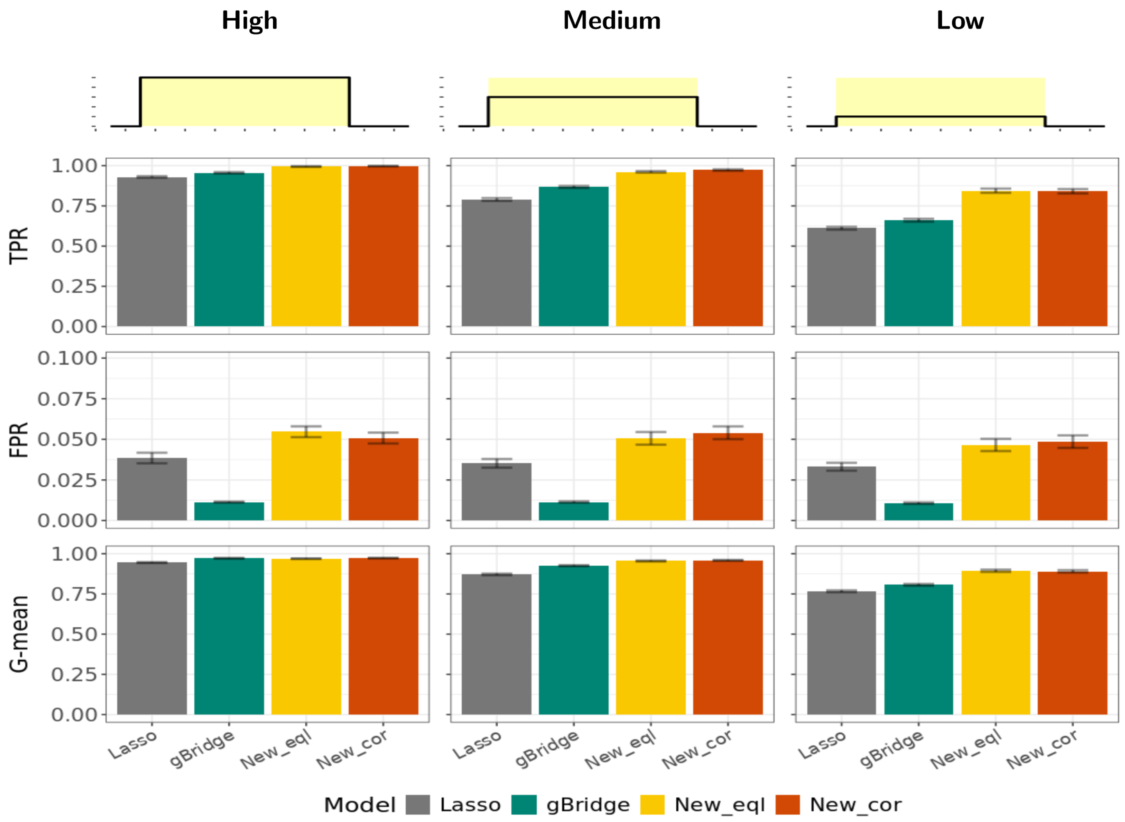


**Figure S4: Variable selection performance evaluated by TPR, FPR, and G-mean under Simulation Scenario 2 for continuous traits.** The top row illustrates the design of causal signals under Scenario 2, with three levels of signal strength: high (left column), medium (middle column) to low (right column). The second to fourth rows respectively show the true positive rate (TPR), the false positive rate (FPR), and the geometric mean (G-mean) of TPR and (1-FPR). All metrics are reported as the mean values of 100 simulation replications, with bars indicating their standard errors.

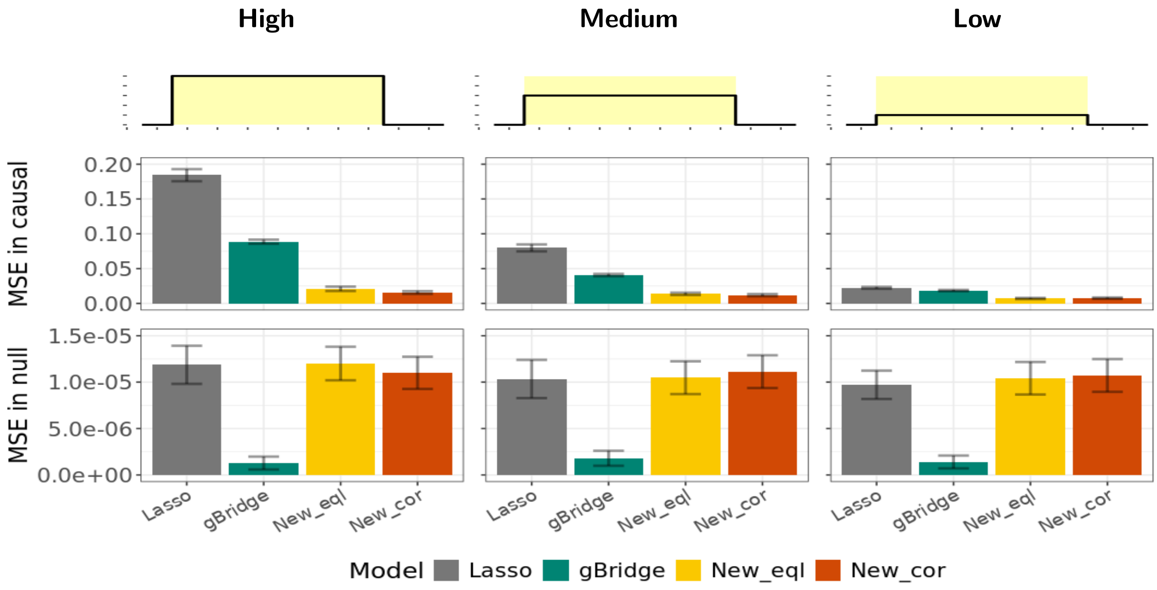


**Figure S5: Effect estimation performance evaluated by mean squared error (MSE) under Simulation Scenario 2 for continuous traits.** The top row illustrates the design of causal signals under Scenario 2, with three levels of signal strength: high (left column), medium (middle column) to low (right column). The second and third rows show the MSEs in causal region and null (non-causal) regions, respectively. MSEs are reported as the mean values of 100 simulation replications, with bars indicating their standard errors.

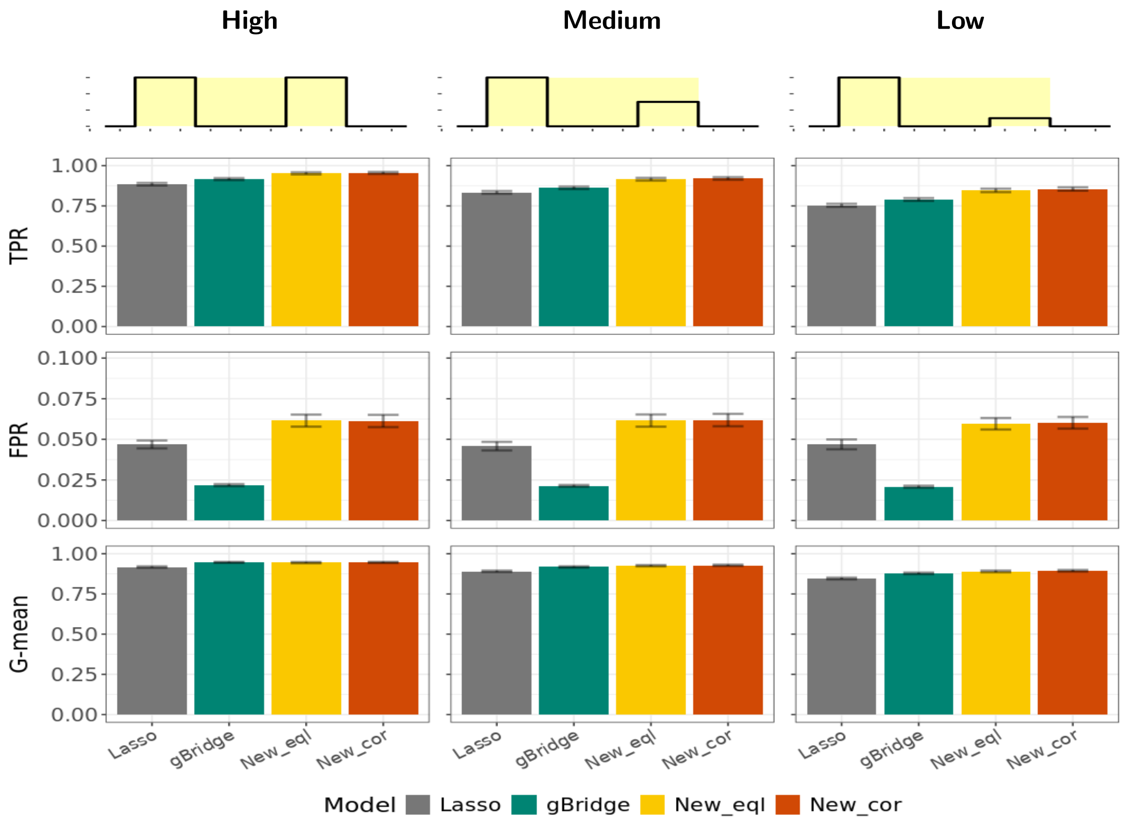


**Figure S6: Variable selection performance evaluated by TPR, FPR, and G-mean under Simulation Scenario 3 for continuous traits.** The top row illustrates the design of causal signals under Scenario 3, with three levels of signal strength: high (left column), medium (middle column) to low (right column). The second to fourth rows respectively show the true positive rate (TPR), the false positive rate (FPR), and the geometric mean (G-mean) of TPR and (1-FPR). All metrics are reported as the mean values of 100 simulation replications, with bars indicating their standard errors.

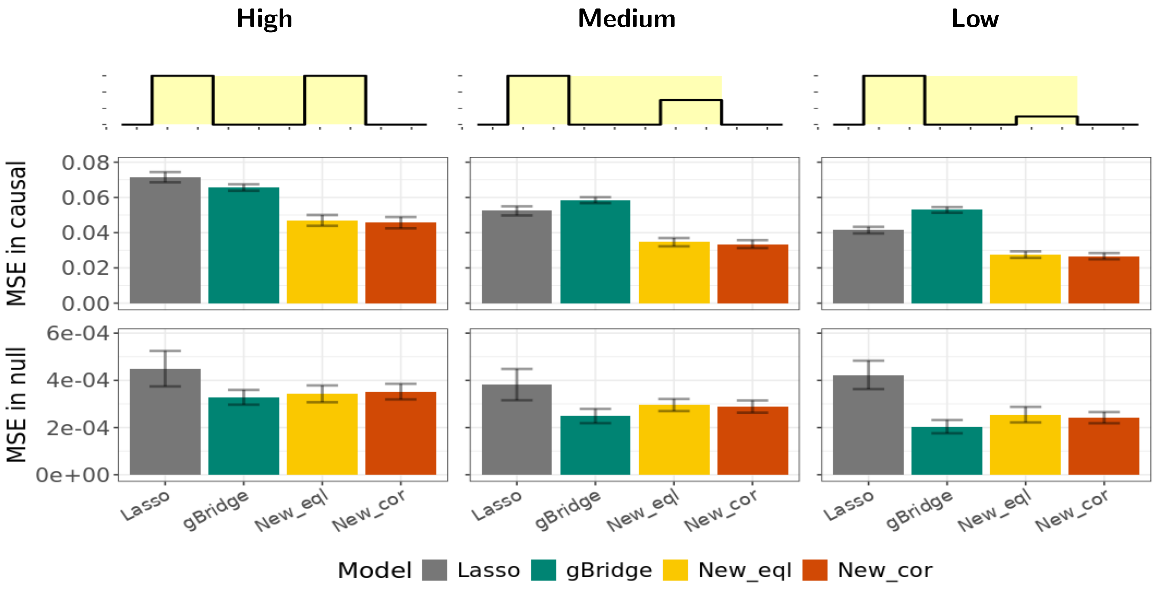


**Figure S7: Effect estimation performance evaluated by mean squared error (MSE) under Simulation Scenario 3 for continuous traits.** The top row illustrates the design of causal signals under Scenario 3, with three levels of signal strength: high (left column), medium (middle column) to low (right column). The second and third rows show the MSEs in causal region and null (non-causal) regions, respectively. MSEs are reported as the mean values of 100 simulation replications, with bars indicating their standard errors.
